# Protocadherin 9 promotes cell survival of different bipolar subtypes in the developing mouse retina

**DOI:** 10.64898/2026.04.17.719213

**Authors:** Marlon F Mattos, Daniela Becerril, Jingyao Guo, Cesiah C Gomez, Elizabeth Zuniga-Sanchez

**Author notes:** **Competing Financial Interests.** The authors declare no competing financial interests.

## Abstract

Neural circuit assembly relies on different neuronal subtypes coming together to form a functional circuit. The question of how the appropriate number of each subtype is integrated into an emerging circuit remains relatively unknown. To answer this question, we used the mouse retina to uncover the molecular mechanisms responsible for neuron subtype integration in a developing circuit. In the mammalian retina, bipolar neurons are a class of interneurons that relay visual information from photoreceptors to ganglion cells. Extensive studies have shown there are 15 distinct bipolar subtypes: 6 types of OFF cone bipolars, 8 types of ON cone bipolars, and 1 type of rod bipolar. During retinal development, bipolar neurons are born in excess and through programmed cell death, a precise number of each subtype remains to give rise to the retinal circuit. Although this process has been well-described, little is known about the key molecules responsible for bipolar subtype integration in the developing retina. Our work uncovered a new role for the autism-associated risk gene, Protocadherin 9 (Pcdh9) in bipolar subtype integration. Deletion of Pcdh9 using a floxed allele leads to loss of OFF and ON cone bipolars; however, disruption in the extracellular binding of Pcdh9 leads to selective loss of ON cone bipolars but not rod bipolars. Moreover, we found this later function of Pcdh9 is mediated by homophilic interactions between ON cone bipolars and their known synaptic partners. Taken together, our work revealed a new role for Pcdh9 in bipolar subtype integration during retinal development.

**SUMMARY STATEMENT:** Neural circuits are comprised of multiple neuronal subtypes where a specific number need to come together to give rise to a functional circuit. Although this is a critical process during neurodevelopment, little is known about the molecular mechanisms that determines the precise number of each subtype during circuit development. In the present study, we identified the autism risk gene, Protocadherin 9 as a critical molecule in subtype integration of bipolar neurons within the developing mouse retina. Using newly generated mouse lines, we found distinct requirements of Pcdh9 to promote survival in different bipolar subtypes during retinal circuit assembly. The significance of this work is that it shed lights into how different neuronal subtypes are integrated in nascent neural circuits.

## INTRODUCTION

A functional neural circuit is comprised of multiple neuronal subtypes where a specific number come together during circuit development. The molecular mechanism that determines the precise number of each subtype in an emerging neural circuit is relatively unknown. To tackle this fundamental question, we used the mouse retina as a model system to identify the key molecules that ensure the appropriate number of neuronal subtypes during retinal circuit assembly. In the mammalian retina, bipolar neurons are a class of interneurons that relay visual information from photoreceptors to ganglion cells ^1^. Bipolar neurons are one of the most well-characterized cell types of the mouse retina with extensive studies documenting their development ^2,3^, gene expression profiles ^4–6^, connectivity ^7,8^, and functional properties ^9,10^. Based on these studies, there are 15 well-characterized types of bipolar neurons: 6 OFF cone bipolars, 8 ON cone bipolars, and 1 rod bipolar. **See Figure 1A**. In the outer retina, dendrites of bipolar neurons synapse selectively to distinct classes of photoreceptors and horizontal cells in the outer plexiform layer (OPL). Cone bipolars mainly synapse to cone photoreceptors and the dendrites of horizontal cells, whereas rod bipolars synapse to rod photoreceptors and the axon terminal of horizontal cells ^4,11,12^. Similarly, within the inner retina, axon terminals of bipolar neurons terminate at distinct layers in the inner plexiform layer (IPL) where they synapse selectively to distinct classes of amacrine cells and ganglion cells ^1^. See **Figure 1A**. The distinct wiring patterns and unique expression profiles of the different bipolar subtypes are important to give rise to the OFF and ON visual pathways of the mammalian retina. However, how these different bipolar subtypes become integrated into the developing retinal circuit remains poorly understood.

**Figure 1:**
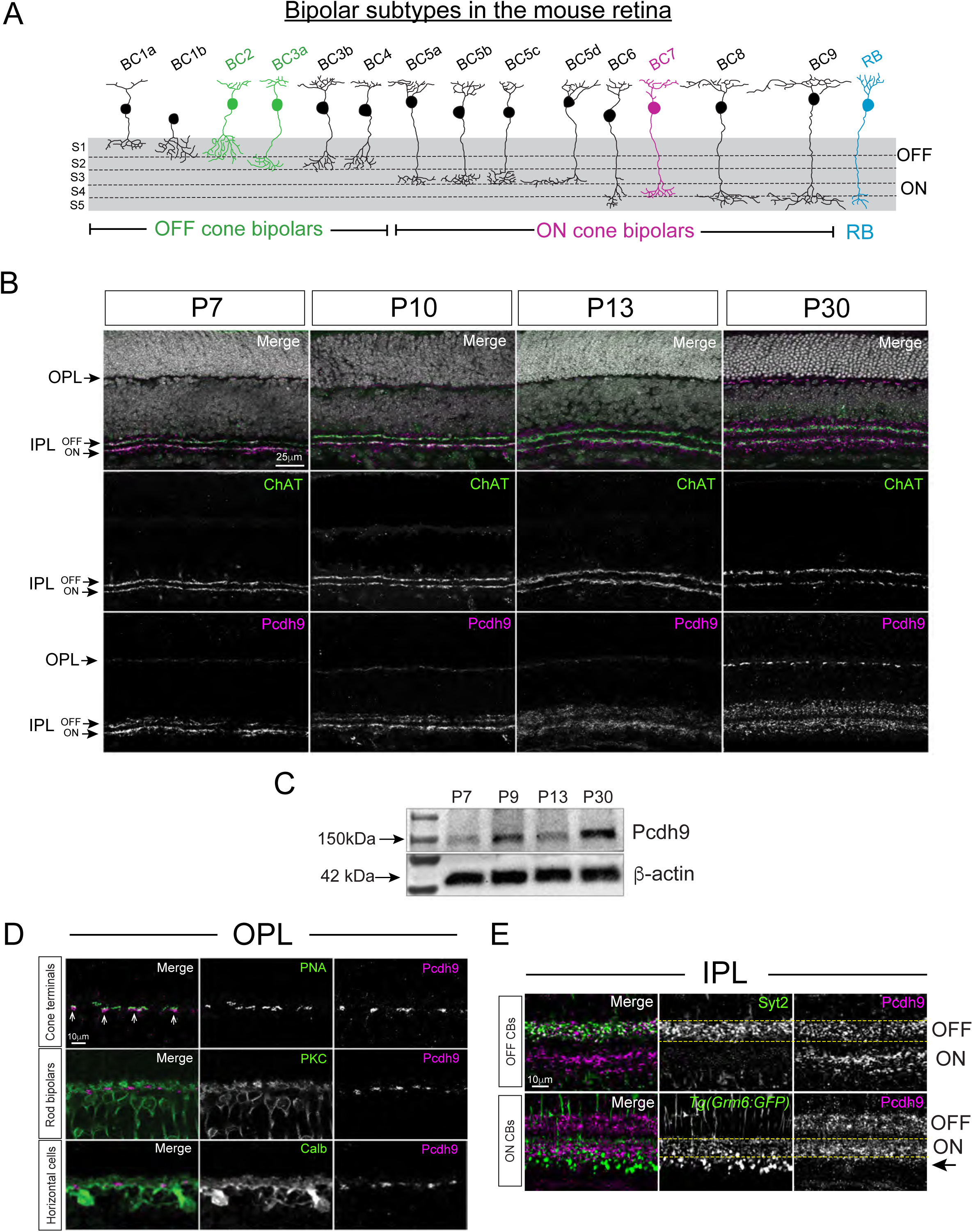
Pcdh9 is localized to the developing synaptic layers in the mouse retina (A-E). Schematic drawing of the 15 different bipolar subtypes in the mouse retina (A). Image adapted from Shekhar et al., 2016. Pcdh9 protein expression (magenta) is detected at P7 in the emerging IPL and then in the OPL at later stages of wild-type retinas (B). Pcdh9 (magenta) co-localizes with anti-Chat (green) which labels the OFF and ON layers of the IPL in (B). Nuclei is stained with DAPI (white) in (B). Western blot analysis at different developmental time points confirms Pcdh9 protein increases from P7 to P30 in wild-type retinas (C). In the OPL, Pcdh9 protein (magenta) is detected immediately below cone terminals labeled with PNA (green) at P30 in wild-type retinas as depicted by white arrows in (D). Pcdh9 protein (magenta) does not co-localize with the dendrites of rod bipolars (anti-PKC, green) but is found within processes of horizontal cells (anti-Calb, green) (D). In the IPL, Pcdh9 protein (magenta) is found in both the OFF and ON layers of the IPL where cone bipolars extend their axons to innervate their synaptic partners (E). The axon terminals of OFF cone bipolar subtype 2 (BC2) are stained with anti-Syt2 (green), and ON bipolars which includes ON cone bipolars and rod bipolars are labeled in the *Tg(Grm6:GFP)* mouse line (green) in wild-type retinas at P30 in (E). Pcdh9 protein is not detected in the S5 layer of the IPL marked by black arrow where the axon terminals of rod bipolars reside (E). Scale bars are shown for each figure.

During development, bipolar neurons are the last neuron type to be born and the last one to become incorporated into the retinal circuit ^13^. Bipolar neurons are born starting at postnatal day (P)0; however, they do not begin to extend their neuronal processes until P5 ^2,13^. The dendrites and axon of bipolar neurons emerge from a long neuro-epithelial process that spans the entire retina, and they extend neurites laterally only in regions of the emerging synaptic layers (i.e. OPL and IPL) ^2^. Next, bipolar neurons undergo a period of programmed cell death from P8-12 where different bipolar subtypes die through cell apoptosis at different frequencies ^14,15^. Research by Keeley and colleagues showed a precise number of bipolars are eliminated through programmed cell death ^14^. This includes 24% of rod bipolars (RB), 39% of cone bipolar type 2 (BC2), 33% of cone bipolar type 3B (BC3B), and 13% of cone bipolar type 4 (BC4) ^14^. Following programmed cell death there is a period of synaptic refinement. Synaptic refinement refers to the process of how dendrites of bipolars undergo remodeling to achieve tiling known as minimal overall between neighbors belonging to the same subtype, and proper innervation to their respective cone photoreceptor target ^8,16^. By P30, bipolar subtype integration and circuit formation in the mouse retina is largely complete.

Although bipolar synapse development has been well-described (reviewed in ^17^), little is known about the molecular mechanisms that ensure how the appropriate number of each bipolar subtype is integrated during retinal development. To identify potential candidates, we searched published RNA sequencing data of bipolars in the mouse retina ^4–6^. We specifically looked for genes that were differentially expressed among the different bipolar subtypes. Our analysis revealed different members of the non-clustered Protocadherin family were differentially expressed among the distinct bipolar subtypes (data not shown). One of these family members is Protocadherin 9 (Pcdh9), which is a known risk gene for autism spectrum disorders ^18,19^, and shown to be important in synapse formation of the developing mouse hippocampus ^20^. Even though Pcdh9 has been described as a critical molecule for normal brain development, little is known about its function in the developing retina. Thus, we set out to investigate if Pcdh9 may be responsible for bipolar subtype integration during retinal circuit development.

## RESULTS

### Pcdh9 is expressed in the developing OPL and IPL of the mouse retina

We first examined Pcdh9 protein expression by performing antibody staining in wild-type retinas at different developmental time points. See **Figure 1B**. Our data revealed Pcdh9 is expressed in the IPL and co-localizes with the amacrine cell marker, anti-ChAT in both the OFF and ON layers of the IPL at P7 (**Figure 1B**). Pcdh9 expression in the IPL continues from P10 to P30, with more broad expression seen at later stages when the IPL is fully mature (**Figure 1B**). We also detected Pcdh9 expression in the OPL starting at P10 and continues to P30 (**Figure 1B**). Western blots of Pcdh9 protein expression in wild-type retinas confirms increasing expression from P7 to P30 as shown in **Figure 1C**. Close examination of Pcdh9 expression in the synaptic layers (OPL, IPL) revealed Pcdh9 is localized to the base of cone terminals stained with PNA as depicted by white arrows in **Figure 1D**. We also noticed Pcdh9 protein co-localizes with the processes of horizontal cells (anti-Calb) but not with the dendrites of rod bipolars (anti-PKC) as shown in **Figure 1D**. In the IPL, Pcdh9 protein (magenta) is found in both the OFF and ON sublamina and co-localizes with anti-Syt2 which labels the axon terminals of a subtype of OFF cone bipolars, and GFP from the *Tg(Grm6:GFP)* mouse line which labels ON cone bipolars and rod bipolars (**Figure 1E**). However, Pcdh9 does not appear to express in the sublamina of the IPL where rod bipolars reside as shown by black arrow in **Figure 1E**. These data show Pcdh9 is localized to the developing synaptic layers where the dendrites and axon terminals of cone bipolars reside.

### New mouse lines to investigate Pcdh9 function in synapse development

As Pcdh9 germline mutants are known to die postnatally ^21^, we generated new tools to study the function of Pcdh9 at later developmental time points. This includes a floxed allele of Pcdh9 that is designed to excise Exon 2 via cre-mediated recombination as illustrated in **Supplementary Figure 1A**. We crossed our floxed allele of Pcdh9 to the *Tg(Chx10-cre)* mouse line ^22^, to conditionally delete Pcdh9 throughout the retina (referred to as Pcdh9 CKO) as shown in **Figure 2A**. We also generated a transgenic mouse that harbors a single point mutation (L385D) in the extracellular domain of Pcdh9 known to inhibit cell-cell interactions based on published biophysical data ^23^. See **Figure 2A** and **Supplementary Figure 2A**. These mice were bred to be homozygous for the L385D point mutation and referred to as Pcdh9^L385D^. Using these new mouse lines, we first confirmed loss of Pcdh9 in Pcdh9 CKO retinas by performing *in situ* hybridization and antibody staining. We used an mRNA probe that recognizes Exon 2 of the *Pcdh9* gene as shown in **Supplementary Figure 1A** to confirm excision within the retina. As expected, we found reduced *Pcdh9* mRNA expression in the nuclear layers (INL, GCL) of Pcdh9 CKO compared to controls at P30 (**Figure 2B,C**). Consequently, Pcdh9 protein expression is also significantly reduced in the OPL and IPL of Pcdh9 CKO retinas (shown in green) compared to controls (**Figure 2D,E**). Next, we compared the expression levels of Pcdh9 in Pcdh9^L385D^ animals with antibody staining and *in situ* hybridization. Our data show no significant difference in *Pcdh9* mRNA expression within the INL and GCL of Pcdh9^L385D^ animals and controls (**Figure 2B,C**). Furthermore, we see no significant difference in anti-Pcdh9 fluorescence intensity across the different retinal layers of Pcdh9^L385D^ (shown in magenta) compared to controls (**Figure 2D,E**). These results demonstrate that we can efficiently knockdown Pcdh9 in the retina of Pcdh9 CKO animals, with minimal disruption of Pcdh9 expression in Pcdh9^L385D^ transgenic mice.

**Figure 2:**
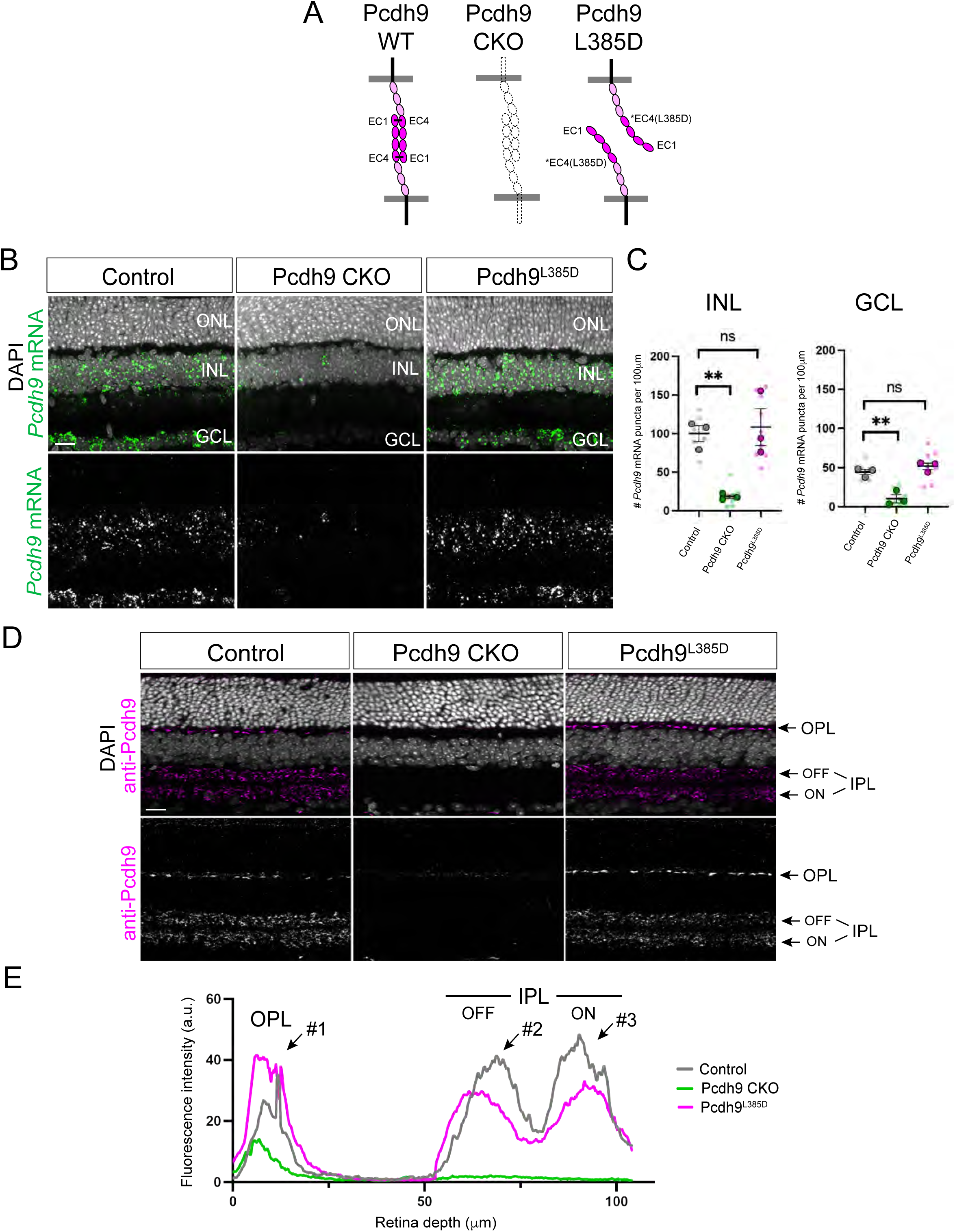
Generation of new mouse lines to study Pcdh9 in retinal development (A-E). Schematic representation of how the extracellular domain of Pcdh9 participates in homophilic trans (cell-cell) interactions within the different transgenic mouse lines (A). Wild-type Pcdh9 forms dimers in an antiparallel manner via the extracellular cadherin (EC) repeats, EC1 and EC4 as shown in control mice in (A). Pcdh9 CKO lacks full-length Pcdh9 protein and do not form Pcdh9-mediated cell-cell interactions (A). Pcdh9^L385D^ animals express Pcdh9 protein but contain a point mutation (L385D) in EC4 that prevents binding with EC1, abolishing cell-cell binding (A). *Pcdh9* mRNA (shown in green) was detected via *in situ* hybridization in controls, Pcdh9 CKO, and Pcdh9^L385D^ at P30 in (B). *Pcdh9* mRNA is highly expressed in the INL and GCL but absent in the ONL of wild-type retinas as shown in controls (B). *Pcdh9* mRNA levels are significantly reduced in the INL and GCL of Pcdh9 CKO animals but not in Pcdh9^L385D^ mice (B). DAPI staining was used to visualize the different nuclear layers. Quantification of the total number of *Pcdh9* mRNA puncta (0.6μm in size) per 100μm of OPL length in the INL and GCL is shown in (C). A total of 9 retinal sections (small circles) from three different animals (big circles) were used for quantification. Data are represented as mean values ± SEM with statistical significance determined by an unpaired two-tailed Student’s t test. ns p > 0.05, **p < 0.01. Pcdh9 protein expression (anti-Pcdh9, magenta) is detected in the synaptic layers (i.e. OPL, IPL) in wild-type retinas at P30 as shown in controls (D). Pcdh9 protein expression is significantly reduced throughout the retina in Pcdh9 CKO animals compared to controls but not affected in Pcdh9^L385D^ retinas (D). Measurements of the anti-Pcdh9 fluorescence intensity across the retina in controls, Pcdh9 CKO, and Pcdh9^L385D^ are shown in (E). Three distinct peaks can be seen where Pcdh9 fluorescence intensity is the highest as depicted by black arrows in (E). This includes the OPL (#1), OFF layer of the IPL (#2), and the ON layer of the IPL (#3). All three peaks are reduced in Pcdh9 CKO (green) compared to controls (grey) but not in Pcdh9^L385D^ (magenta) as shown in (E). Scale bar = 25μm.

### Differences in OFF cone bipolar survival between Pcdh9 CKO and Pcdh9^L385D^ animals

Using our newly generated mouse lines, we then examined the different bipolar subtypes at P30 when synapse formation in the mouse retina is complete. By P30, bipolar subtypes could be easily identified using available antibodies and mouse lines that capture their distinct gene expression and morphology ^1,5^. To investigate the function of Pcdh9 in OFF cone bipolars, we used the Syt2 antibody that has been shown to label cone bipolar type 2 (BC2) and anti-HCN4 for cone bipolar type 3a (BC3a) ^1^. See **Figure 1A**. At P30, the axon terminals of wild-type BC2 (shown in green) are confined to the S1-S2 of the OFF layer in the IPL as seen in controls in **Figure 3A**. Loss of Pcdh9 results in several gaps of Syt2 staining within the S1-S2 layer of the IPL in Pcdh9 CKO animals (**Figure 3A**). However, these gaps in Syt2 staining are not seen in Pcdh9^L385D^ retinas (**Figure 3A**). To quantify these observations, we used the published *IPLaminator* tool to measure fluorescence intensity across the different layers of the IPL ^24^. Using the *IPLaminator* tool, we found a reduction of Syt2 staining (shown in green) within the S1-S2 layer of the IPL in Pcdh9 CKO but not in Pcdh9^L385D^ compared to controls as depicted by black arrow in **Figure 3B**. The amacrine cell marker, anti-ChAT labels S2 and S4 layer of the IPL and was used as a reference in **Figure 3A,B**. Moreover, these gaps in Syt2 staining in Pcdh9 CKO were due to loss of BC2 neurons as wholemount retinal images stained with anti-Syt2 showed fewer cell bodies in regions where there are no axon terminals (**Figure 3C**). Quantification of Syt2-positive cell bodies from wholemount retinal images from controls, Pcdh9 CKO, and Pcdh9^L385D^ animals are shown in **Figure 3D**. We also examined cone bipolar type 3a (BC3a) using the HCN4 antibody, and found a similar loss of BC3a protein expression within the S2 layer of the IPL in Pcdh9 CKO but not in Pcdh9^L385D^ animals compared to controls (**Figure 3E**). These findings were also quantified across different animals by measuring the fluorescence intensity across the IPL as shown in **Figure 3F**. Our results demonstrate conditional deletion of Pcdh9 or Pcdh9 CKO affects survival of BC2 and BC3a neurons, whereas disruption in the extracellular binding of Pcdh9 in Pcdh9^L385D^ animals does not affect these OFF cone bipolar subtypes.

**Figure 3:**
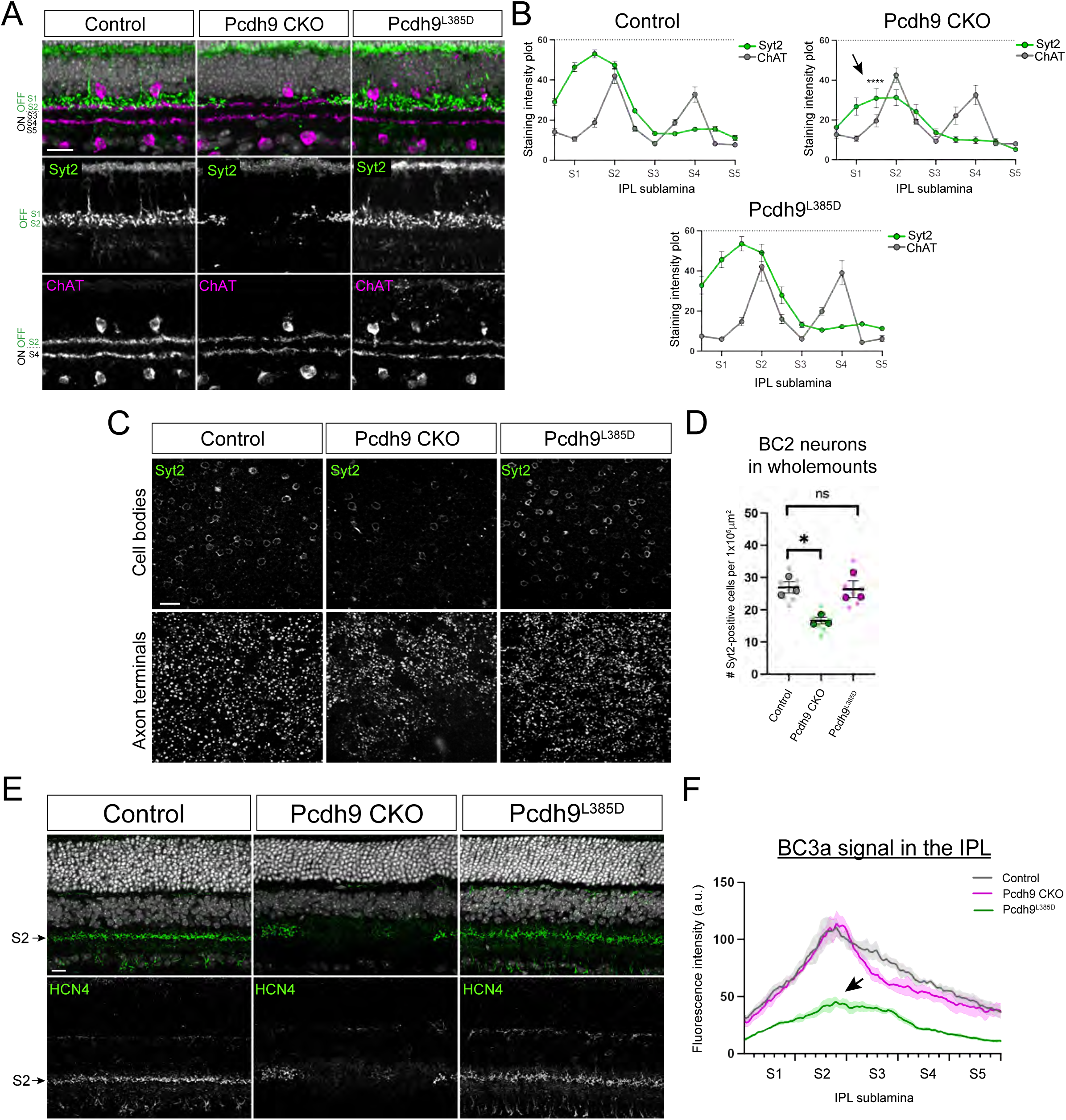
Deletion of full-length Pcdh9 but not disruption in extracellular binding leads to loss of OFF cone bipolars (A-F). At P30, the axon terminals of BC2 neurons labeled with anti-Syt2 (green) reside in the OFF S1-S2 layer of the IPL, and form a continuous layer of Syt2 expression in wild-type retinas as shown in controls (A). Deletion of full-length Pcdh9 or Pcdh9 CKO animals but not Pcdh9^L385D^ results in gaps or areas devoid of Syt2 staining shown in green compared to controls (A). ChAT staining (magenta) which labels amacrine cells was used as a reference to label the OFF (S2) and ON (S4) layers of the IPL in (A). Nuclei was stained with DAPI (A). Fluorescence intensity of anti-Syt2 (green) and anti-ChAT (grey) in the IPL was measured in the different transgenic mouse lines using the *IPLaminator* tool (B). A total of 9 retinal sections from three different animals were used for quantification. Syt2 staining in the S1-S2 layer was decreased in Pcdh9 CKO (black arrow) but not in Pcdh9^L385D^ compared to controls (B). ChAT staining was consistent across all three groups (B). Data are represented as mean values ± SEM. Measurements within the S1-S2 layer of the IPL of Pcdh9 CKO retinas were statistically significant compared to controls as determined by a Holm–Sidak method for multiple comparisons with ****p < 0.0001. Wholemount retina images confirm reduced Syt2 staining in the IPL of Pcdh9 CKO are due to loss of BC2 neurons (C). The number of Syt2-positive cell bodies per normalized area of wholemount retinas in the different groups are shown in (D). Cell counts were performed on three different regions of the retina (small circles) from three different animals (big circles) per group in (D). Data are represented as mean values ± SEM. Statistical significance determined by an unpaired two-tailed Student’s t test. ns p > 0.05, *p < 0.05. BC3a were labeled with anti-HCN4 (green) and showed similar gaps in staining within the S2 layer of the IPL in Pcdh9 CKO animals but not in controls or Pcdh9^L385D^ retinas (E). Nuclei were stained with DAPI in (C). Fluorescence intensity of HCN4 staining within the IPL were measured in all three groups in (F). Reduced HCN4 staining in the S2 layer was observed in Pcdh9 CKO (green line) as depicted by black arrow but not in Pcdh9^L385D^ (magenta) or controls (grey) (F). Measurements were performed using three different retinal sections from three different animals per group and shown as mean values ± SEM (E). Scale bar = 25μm.

### Loss of ON cone bipolars but not rod bipolars in retinas with disruption of Pcdh9

Next, we examined the role of Pcdh9 in ON bipolars using the *Tg(Gus:GFP)* mouse line, which has been shown previously to label cone bipolar type 7 (BC7) and rod bipolars ^1,16,25^. We crossed *Tg(Gus:GFP)* mice and Pcdh9^L385D^ animals and stained the retinas with the known rod bipolar marker, anti-PKC to distinguish between BC7 and rod bipolars. Cells that were GFP-positive and PKC-negative were considered BC7 neurons, whereas those that were GFP-positive and PKC-positive were rod bipolars. Retinal wholemounts of Pcdh9^L385D^ animals showed a significant loss of BC7 neurons (shown in green) but not rod bipolars (shown in magenta) when compared to controls (**Figure 4A,B**). However, in the case of Pcdh9 CKO animals, the *Tg(Chx10-cre)* mouse line contains a transgene consisting of a fusion protein between Cre recombinase and eGFP, which results in green fluorescence expression in the cell bodies of bipolars as previously reported ^22^. This made it difficult to distinguish between BC7 neurons and rod bipolars in retinal wholemounts of Pcdh9 CKO animals crossed with the *Tg(Gus:GFP)* reporter. Despite the GFP signal from *Tg(Chx10-cre)* mouse line, this expression was largely confined to the cell bodies and not the axon terminals of bipolar neurons. Thereby, we were able to analyze the axon terminals of BC7 neurons and rod bipolars in Pcdh9 CKO and Pcdh9^L385D^ animals crossed with *Tg(Gus:GFP)* as shown in **Figure 4C**. Our data shows similar gaps in the S4 layer of the IPL where the axon terminals of BC7 neurons reside in both Pcdh9 CKO and Pcdh9^L385D^ animals compared to controls (**Figure 4C**). However, no gaps were observed in the S5 layer of the IPL where the axon terminals of rod bipolars labeled with anti-PKC (magenta) are confined (**Figure 4C**). These findings were quantified across multiple animals using the *IPLaminator* tool and shown in **Figure 4D,E**. Our results confirm there is a loss of *Tg(Gus:GFP)* intensity (shown in cyan) within the S4 layer of the IPL in Pcdh9 CKO and Pcdh9^L385D^ compared to controls as depicted by black arrows in **Figure 4D**. However, no significant difference was observed for PKC (magenta) or ChAT (grey) between controls, Pcdh9 CKO, and Pcdh9^L385D^ animals (**Figure 4E**). To further validate the loss of BC7 neurons in Pcdh9 CKO and Pcdh9^L385D^ animals, we performed *in situ* hybridization using the known BC7 marker, *Igfn1* that has been described previously ^5^. We found *Igfn1* mRNA is expressed in evenly spaced clusters (shown by yellow dotted circles) that span the entire INL as shown in controls in **Supplemental Figure 3A**. Disruption of Pcdh9 function leads to loss of *Igfn1*-positive cell clusters in Pcdh9 CKO and Pcdh9^L385D^ compared to controls, which is consistent with loss of BC7 neurons (**Supplemental Figure 3A,B**). These results demonstrate full-length deletion of Pcdh9 leads to loss of BC2, BC3a, and BC7 but not rod bipolars. However, disruption in the extracellular binding of Pcdh9 or Pcdh9^L385D^ results in selective loss of BC7 but not BC2, BC3a, or rod bipolars.

**Figure 4:**
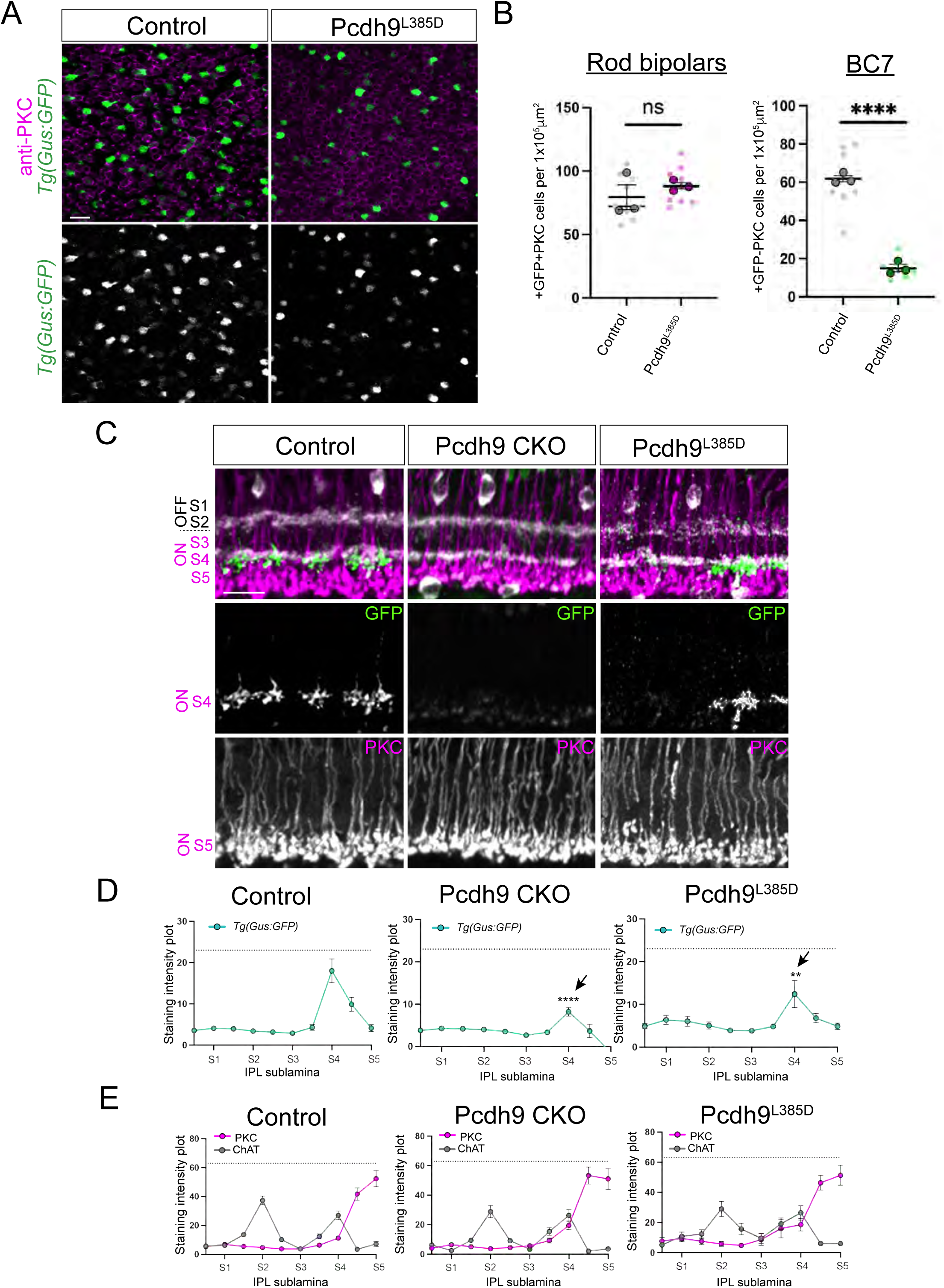
Disruption of Pcdh9 function results in loss of ON cone bipolars but not rod bipolars (A-E). Wholemount retinal images of Pcdh9^L385D^ animals crossed with *Tg(Gus:GFP)* and stained with anti-PKC to label BC7 neurons and rod bipolars in (A). Controls are *Tg(Gus:GFP)* that do not harbor the L385D point mutation in the extracellular domain of Pcdh9. BC7 neurons are GFP-positive and PKC-negative, whereas rod bipolars are GFP-positive and PKC-positive in (A). BC7 neurons are reduced but not rod bipolars in Pcdh9^L385D^ compared to controls (A). Cell counts of BC7 neurons and rod bipolars per normalized area in wholemount images are shown in (B). Data are represented as mean values ± SEM. Three different measurements from wholemount images (small circles) from three different animals (big circles) were used for quantification. Statistical significance was determined by an unpaired two-tailed Student’s t test. ns p > 0.05, ****p < 0.0001. Retinal sections from Pcdh9 CKO and Pcdh9^L385D^ animals crossed with *Tg(Gus:GFP)* and stained with anti-PKC and anti-ChAT are shown in (C). The axon terminals of BC7 neurons (GFP, green) are evenly spaced in the S4 layer of the IPL, whereas the axon terminals of rod bipolars (PKC) are confined to the S5 layer in wild-type retinas as shown in controls (C). Significant gaps in labeling of the axon terminals of BC7 neurons (shown in green) are observed in Pcdh9 CKO and Pcdh9^L385D^ retinas but not in the axon terminals of rod bipolars (shown in magenta) (C). Measurements of the staining intensity across the different layers of the IPL in (C) using the *IPLaminator* tool are shown in (D,E). The GFP signal in S4 is reduced in both Pcdh9 CKO and Pcdh9^L385D^ retinas depicted by black arrows compared to controls (D). However, PKC staining is similar across all transgenic mouse lines showing a high peak of expression at S5 (E). ChAT staining (amacrine cell marker) shown in grey was used a reference to label S2 and S4 layer of the IPL in (E). A total of 9 retinal sections from three different animals per group were used for quantification. Data are shown as mean values ± SEM with statistical significance determined by Holm–Sidak method for multiple comparisons. **p < 0.01, ****p < 0.0001. Scale bar = 25μm.

### Distinct changes in postsynaptic protein expression in animals with altered Pcdh9 function

We found disruption of Pcdh9 function leads to loss of different subtypes of cone bipolars. To further confirm this observation, we examined expression of known post-synaptic receptors known to be selectively expressed in distinct bipolar subtypes ^5^. This includes the post-synaptic kainate receptor GluK1 known to be expressed on the dendrites of OFF cone bipolars ^5,26,27^, and the metabotropic Glutamate receptor 6 (mGluR6) which is found on the dendrites of ON cone bipolars and rod bipolars ^5,28^. At P30, mGluR6 protein expression is localized to the dendrites of both rod bipolars and ON cone bipolars but adopt a distinct morphology at rod and cone synapses ^29^. At rod synapses, mGluR6 appears as a small, round puncta-like structure, whereas at cone synapses, multiple mGluR6 cluster together and form a straight line as seen in controls in **Figure 5A**. Moreover, we can isolate mGluR6 protein expression within cone synapses using PNA staining as shown by dotted circles in **Figure 5A**. We found mGluR6 within cone synapses is significantly reduced in Pcdh9 CKO and Pcdh9^L385D^ retinas compared to controls (**Figure 5A**). To quantify this observation, we measured the relative fluorescence signal of mGluR6 across different cone synapses as described previously ^26^. Indeed, we found reduced mGluR6 protein expression (magenta) in both Pcdh9 CKO and Pcdh9^L385D^ (black arrows) compared to controls, with minimal change in PNA staining shown in green (**Figure 5A**). Next, we assessed GluK1 protein expression at cone synapses, which is known to be selectively expressed in the dendrites of OFF cone bipolars ^5,26^. For this analysis, we used the pre-synaptic marker, anti-CtBP2 shown in magenta to identify cone terminals in **Figure 5B**. Similar to mGluR6, GluK1 protein at P30 appears as a straight line localized to the base of cone terminals as shown in controls in **Figure 5B**. We measured the relative fluorescence signal of GluK1 expression in different cone synapses of controls, Pcdh9 CKO, and Pcdh9^L385D^ retinas (**Figure 5B**). Our results show a significant reduction of GluK1 fluorescence signal (green line) in Pcdh9 CKO as depicted by black arrow but not in Pcdh9^L385D^ retinas compared to controls (**Figure 5B**). These data demonstrate a reduction of mGluR6 within synapses of ON cone bipolars in both Pcdh9 CKO and Pcdh9^L385D^ retinas, and only a decrease in GluK1 of OFF cone bipolars in Pcdh9 CKO animals. In addition, we counted the number of cone terminals found within the OPL and examined protein expression of known pre-synaptic markers such as Bassoon. We found no statistical difference in the number of cone terminals or pre-synaptic Bassoon between controls, Pcdh9 CKO, and Pcdh9^L385D^ animals (**Supplementary Figure 4A-C**). These findings suggest loss of mGluR6 or GluK1 protein expression in the OPL cannot be explained by absence of cone photoreceptors. Moreover, our data confirms the selective loss of ON cone bipolars in Pcdh9^L385D^ animals, whereas both OFF and ON cone bipolars are affected due to loss of full-length Pcdh9 or Pcdh9 CKO.

**Figure 5:**
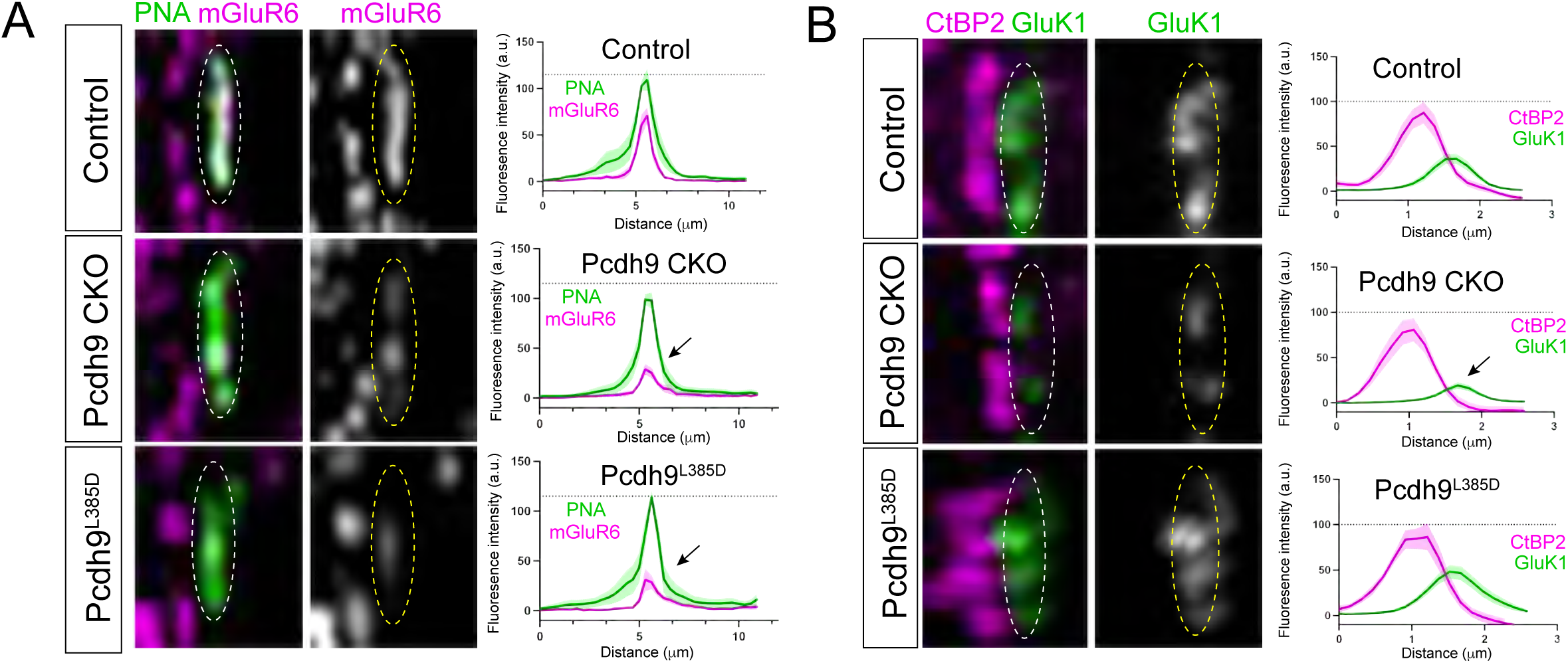
Reduced protein expression of post-synaptic receptors at cones synapses due to loss of Pcdh9 function (A-B). mGluR6 protein expression (magenta) is reduced at cone synapses labeled with PNA (green) in Pcdh9 CKO and Pcdh9^L385D^ retinas compared to controls (A). Measurements of the fluorescence intensity of PNA staining (green) and mGluR6 protein expression (magenta) is shown on the right panel of (A). GluK1 protein expression (green) is also reduced at cone synapses in Pcdh9 CKO as shown by dotted circles but not in Pcdh9^L385D^ compared to controls (B). Cone synapses were identified using the pre-synaptic marker, anti-CtBP2 (magenta) in (B). Fluorescence intensity of GluK1 and CtBP2 signal were measured across different cone synapses as shown in right panel of (B). Reduced GluK1 signal is observed in Pcdh9 CKO (black arrow) but not in Pcdh9^L385D^ retinas (B). A total of 9 cone synapses from three different animals per group were analyzed.

### Developmental analysis of cone bipolar survival in Pcdh9 CKO and Pcdh9^L385D^ retinas

Next, we performed a developmental analysis to address when Pcdh9 function is required for cone bipolar survival. As Pcdh9 is known to influence cell apoptosis in other biological processes ^30^, we assessed the role of Pcdh9 in bipolars during programmed cell death (P8-12). To answer this question, we examined the retinas from Pcdh9 CKO and Pcdh9^L385D^ transgenic animals at three key developmental time points: (i) P7 – beginning of cell death, (ii) P9 – midpoint, and (iii) P13 - end of programmed cell death. We used an antibody against the activated form of caspase-3, which is a known marker of apoptosis in the mouse retina to identify dying cells ^31^, and anti-islet1 staining to identify the location of bipolars within wholemount retinas (**Figure 6A-C**). Our data revealed the number of dying cells were not statistically different between controls, Pcdh9 CKO, and Pcdh9^L385D^ at P7 (**Figure 6A,D**). However, by P9 the number of dying cells decreased in controls compared to Pcdh9 CKO and Pcdh9^L385D^ animals (**Figure 6B,D**). By P13, the number of dying cells decreased across all groups, with a few cells still undergoing cell death in Pcdh9^L385D^ animals at these later stages (**Figure 6C,D**). These findings suggest the full-length of Pcdh9 may be required for cell survival during stages of programmed cell death (P8-12), whereas the extracellular domain of Pcdh9 may have additional roles at later stages (>P13) to promote cone bipolar survival.

**Figure 6:**
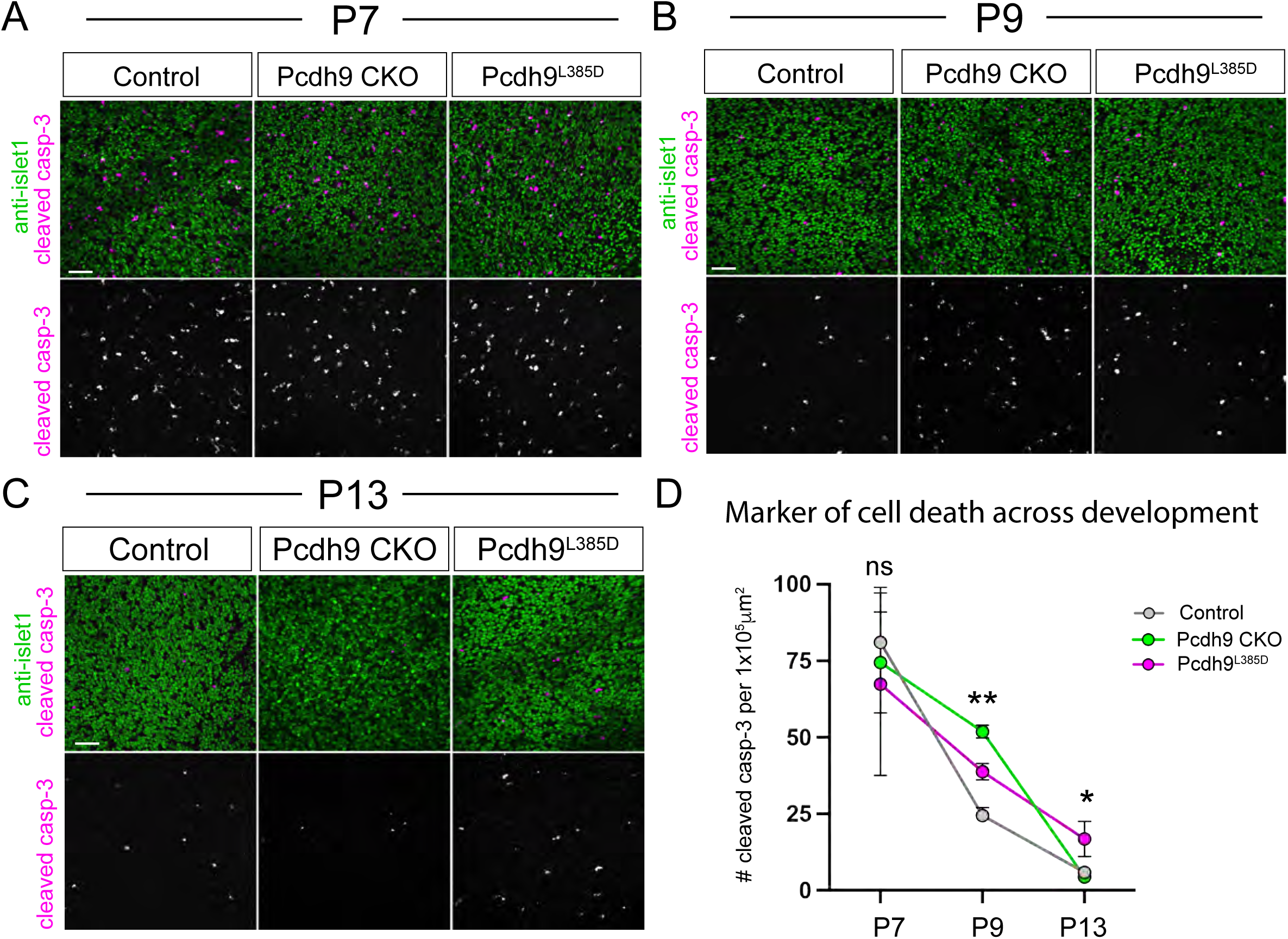
Developmental analysis of cell death in animals with altered Pcdh9 function (A-D). Dying cells were labeled in wholemount retinas with an antibody that recognizes the cleaved form of caspase-3 (cleaved casp-3) shown in magenta at three developmental time points: P7, P9, and P13 in (A-C). Islet1 staining (shown in green) was used to identify the top-most region of the INL (top-INL) where the cell bodies of bipolar neurons reside (A-C). Cell counts of cleaved casp-3-positive cells in the top-INL at the different time points are shown in (D). Three different regions of the retinal wholemounts from three different animals per group were used for quantification in (D). Data are represented as mean values ± SEM with statistical significance determined using an unpaired two-tailed Student’s t-test comparing Pcdh9 CKO and Pcdh9^L385D^ to controls at each developmental time point. ns p > 0.05, **p < 0.01: Pcdh9 CKO vs. control at P9 and Pcdh9^L385D^ vs. control at P9, *p < 0.05: Pcdh9^L385D^ vs. control at P13.

### Pcdh9 is required in horizontal cells and amacrine cells for BC7 survival

To further investigate the function of the extracellular domain of Pcdh9 in ON cone bipolar survival at later stages, we examined the role of Pcdh9 in mediating synapse formation. Dendrites of ON cone bipolars are known to form synapses with horizontal cells and cone photoreceptors in the outer retina, whereas the axon terminal of cone bipolars form synapses with amacrine cells and ganglion cells in the inner retina ^1,7^. See **Figure 7A**. We found Pcdh9 is required for bipolar survival from P7-13, which coincides with the timing of synapse formation in the mouse retina ^4^. Thus, we addressed whether homophilic Pcdh9-Pcdh9 interactions between cone bipolars and their respective partners could be responsible for their survival at later stages. To test this possibility, we deleted Pcdh9 from horizontal cells and amacrine cells using our floxed allele of Pcdh9 (*Pcdh9^flox/flox^*). This was done by crossing *Pcdh9^flox/flox^* mice with the *Tg(Ptf1a-cre)* mouse line ^32^ to generate *Ptf1a-cre;Pcdh9^flox/flox^* transgenic animals. Consistent with published single-cell RNA sequencing data ^6^, we found *Pcdh9* mRNA is highly expressed in the cell bodies of horizontal cells and a subset of amacrine cells labeled with anti-calbindin as shown in controls at P30 (**Figure 7B**). Moreover, *Pcdh9* mRNA levels were significantly reduced within the cell bodies of horizontal cells and amacrine cells in *Tg(Ptf1a-cre;Pcdh9^flox/flox^)* compared to controls but not elsewhere (**Figure 7B**). These observations were quantified by measuring the total number of *Pcdh9* mRNA puncta (∼0.6μm in size) in horizontal cells and amacrine cells from controls and *Tg(Ptf1a-cre;Pcdh9^flox/flox^)*. See **Figure 7C**. Our results confirm we can reliably delete Pcdh9 from horizontal cells and amacrine cells using *Tg(Ptf1a-cre;Pcdh9^flox/flox^)* animals. Next, we examined different bipolar subtype populations in *Tg(Ptf1a-cre;Pcdh9^flox/flox^)* at P30 which is the end of synaptogenesis. Interestingly, we found no change in BC2 neurons stained with anti-Syt2 in *Tg(Ptf1a-cre;Pcdh9^flox/flox^)* compared to controls (**Supplementary Figure 5A,B**). However, we did notice a significant reduction in the number of BC7 neurons in *Tg(Ptf1a-cre;Pcdh9^flox/flox^)* animals crossed with the *Tg(Gus:GFP)* reporter as shown in **Figure 7D,E**. Similar to our previous analysis, cells that were +GFP-PKC were considered BC7 neurons, whereas those that were +GFP+PKC as rod bipolars in **Figure 7E**. We also used the *IPLaminator* tool to quantify the fluorescence intensity in the IPL from the axon terminals of rod bipolars and BC7 neurons (**Figure 7F**). Our data revealed a significant reduction of the GFP signal within the S4 layer of *Tg(Ptf1a-cre;Pcdh9^flox/flox^)* animals compared to controls as depicted by black arrow but not within the S5 layer where the axon terminals of rod bipolars reside (**Figure 7F**). These data suggest we are losing BC7 neurons and not rod bipolars at P30 due to loss of Pcdh9 in horizontal cells and amacrine cells. We then addressed if BC7 neurons in *Tg(Ptf1a-cre;Pcdh9^flox/flox^)* were dying at early time points during synaptogenesis similar to Pcdh9^L385D^ animals. To answer this question, we stained retinal wholemounts with the antibody that recognizes the cleaved form of caspase-3 (cleaved casp-3) and anti-islet1 at P13 (**Supplemental Figure 6A**). We found more dying cells in *Tg(Ptf1a-cre;Pcdh9^flox/flox^)* animals compared to controls (**Supplemental Figure 6B**). This was consistent with our previous data in Figure 6C,D showing a perdurance of dying cells in Pcdh9^L385D^ animals even after programmed cell death. Taken together, our findings suggest homophilic Pcdh9-Pcdh9 interactions between BC7 neurons and horizontal cells and amacrine cells are required for bipolar survival at adult stages. However, deletion of Pcdh9 does not affect BC2 cell numbers suggesting that there must be other mechanisms that ensure their survival at later stages.

**Figure 7:**
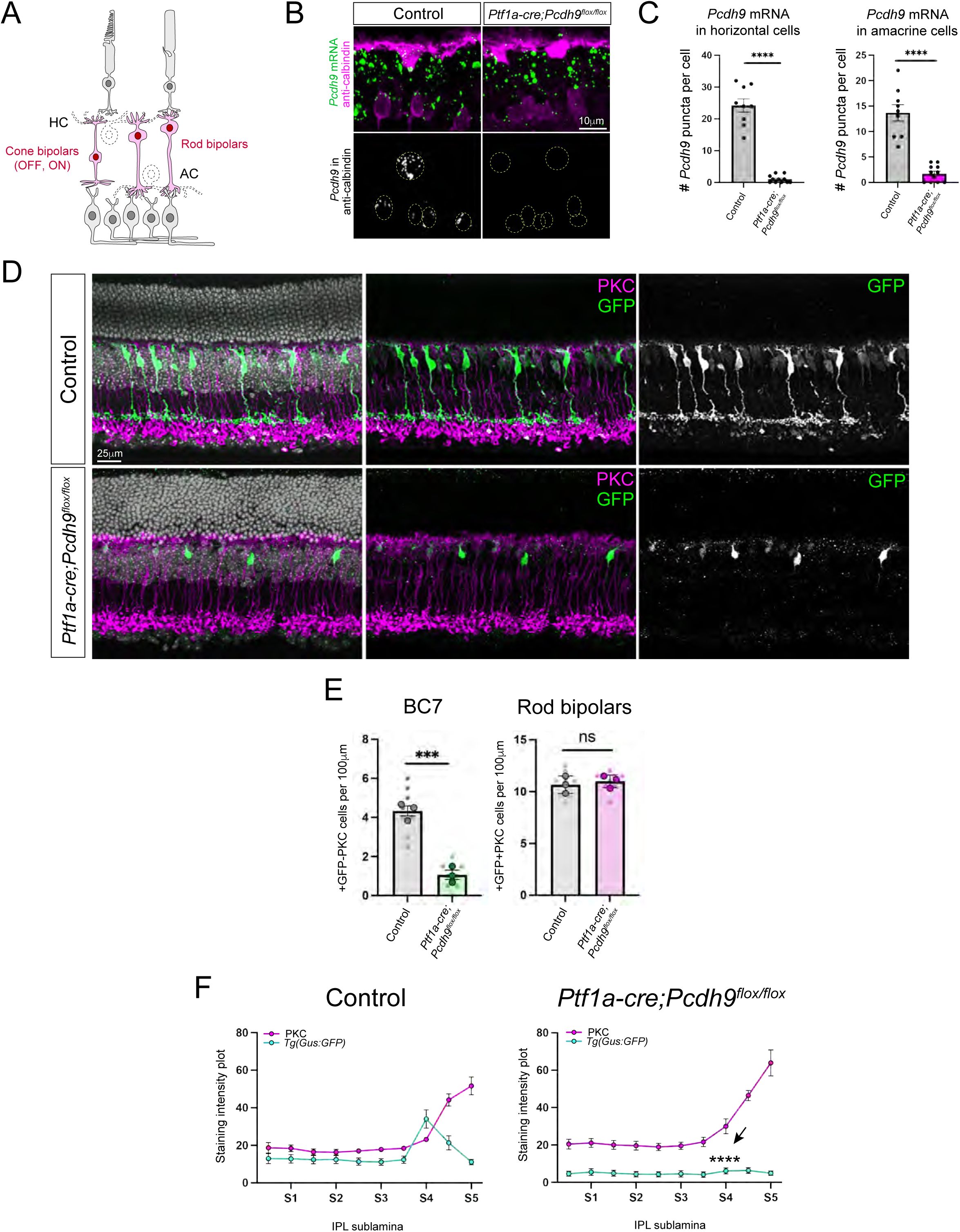
Deletion of Pcdh9 in the synaptic partners of ON cone bipolars leads to loss of BC7 neurons (A-F). Schematic drawing of the mouse retina showing where Pcdh9 was conditionally deleted using *Tg(Ptf1a-cre)* animals (A). Removal of Pcdh9 in horizontal cells and amacrine cells was confirmed via *in situ* hybridization shown in green and antibody staining with calbindin shown in magenta (B). Quantification of the total number of *Pcdh9* mRNA puncta within the cell bodies of horizontal cells and amacrine cells from controls and *Ptf1a-cre;Pcdh9^flox/flox^* animals are shown in (C). Data are represented as mean values ± SEM. A total of n=9 horizontal cells and n=9 amacrine cells from controls and n=11 horizontal cells and n=11 amacrine cells from *Ptf1a-cre;Pcdh9^flox/flox^* were used for analysis. Statistical significance was determined using an unpaired two-tailed Student’s t test. ****p<0.0001. BC7 neurons were labeled by crossing the *Tg(Gus:GFP)* to *Ptf1a-cre;Pcdh9^flox/flox^* animals (referred to as *Ptf1a-cre;Pcdh9^flox/flox^*). Littermates without the cre served as controls. BC7 neurons (green) but not rod bipolars (anti-PKC, magenta) are affected in *Ptf1a-cre;Pcdh9^flox/flox^* animals compared to controls (D). Nuclei with DAPI (white) in (D). The total number of BC7 neurons and rod bipolars in controls and *Ptf1a-cre;Pcdh9^flox/flox^* animals are shown in (E). A total of 9 retinal sections (small circles) from three different animals (big circles) were used for quantification in (E). Data are represented as mean values ± SEM with statistical significance determined using an unpaired two-tailed Student’s t test. ns p>0.05, ***p<0.001. Measurements of the staining intensity using the *IPLaminator* tool for the GFP signal from *Tg(Gus:GFP)* is shown in cyan and PKC staining is shown in magenta in (F). A total of 9 retinal sections from three different animals were used for quantification. Statistical significance was determined by a Holm–Sidak method for multiple comparisons. ****p < 0.0001. Scale bars are shown for each figure.

## DISCUSSION

We uncovered a new role for Pcdh9 in mediating cell survival of distinct bipolar subtype populations in the developing mouse retina. Pcdh9 is expressed at early stages in the nascent OFF and ON layers of the IPL within the inner retina, and then emerges in the synaptic layer or OPL of the outer retina. Conditional deletion of full-length Pcdh9 or Pcdh9 CKO at early developmental time points leads to a selective loss of cone bipolars but not rod bipolars. This includes loss of a subset of OFF and ON cone bipolars while the number of rod bipolars are not affected. Moreover, a single point mutation (L385D) in the extracellular domain of Pcdh9 known to abolish homophilic binding results in similar loss of ON cone bipolars as Pcdh9 CKO. We further demonstrate that this cell loss is due to deletion of Pcdh9 in horizontal cells and amacrine cells which are the known binding partners of ON cone bipolars. These findings highlight an essential role for Pcdh9 in ensuring survival of distinct bipolar subtype populations during retinal development.

### Cadherins in bipolar development

Pcdh9 belongs to the cadherin superfamily of cell adhesion molecules which consists of over 350 members, and they are subdivided into two main classes: the classical and nonclassical cadherins ^33,34^. In the mouse retina, two members of the classical cadherin family, Cadherin-8 (Cdh8) and Cadherin-9 (Cdh9) have been shown to mediate retinal circuit assembly. Specifically, Cdh8 and Cdh9 mediate the positioning of the axon terminals of BC2 and BC5 to the OFF and ON layer within the IPL, respectively ^35^. Global deletion of Cdh8 or Cdh9 results in layer-specific defects of these bipolars and disrupts direction-selective circuits ^35^. However, Duan and colleagues did no report significant cell death of bipolars or other cell types in the absence of Cdh8 and Cdh9 function. This phenotype is different from prior studies reporting loss of members of the nonclassical cadherins which include the clustered Protocadherins. The clustered Protocadherins contain the alpha, beta, and gamma subfamily ^36^, and previous research has shown gamma-Protocadherins are important for cell survival in the developing mouse retina but not for neuronal targeting ^31^. Loss of gamma-Protocadherin leads to cell death of nearly all bipolars including both cone bipolars and rod bipolars ^37^. The widespread effect on all bipolars with loss of gamma-Protocadherin is different from what we found in this study with loss of Pcdh9, a member of the non-clustered Protocadherin family. These findings raise the question whether members of the cadherin family perform specific functions during retinal development, or are they all interchangeable and can compensate for one another? Recent data from the clustered alpha- and gamma-Protocadherin family in the retina suggest that they are not equally redundant with one another ^38^. Loss of alpha-Protocadherins leads to no apparent cell death in the mouse retina, whereas loss of gamma-Protocadherins has a broad effect on survival of multiple retinal cell types ^38^. These results imply that there must be specific requirements of distinct cadherin family members on cell survival during different stages of retinal development.

### Distinct gene expression patterns of non-clustered Protocadherins in bipolars

We propose members of the non-clustered Protocadherin family have distinct requirements on bipolar survival at different stages of retinal circuit formation. This in part is largely based on their temporal and cell-type specific expression. Recent single-cell RNA sequencing data of the mouse retina has revealed that each bipolar subtype expresses a unique set of genes belonging to the non-clustered Protocadherin family ^5,6^. From these published datasets, we found nearly all cone bipolars express Pcdh9 except for rod bipolars. However, most cone bipolar subtypes express at least one or more additional family member. For example, BC1A expresses Pcdh9 and Pcdh17 whereas BC1B expresses Pcdh9 and Pcdh10. Our data suggests OFF cone bipolars may activate separate signaling pathways to promote their survival that do not rely on the extracellular domain of Pcdh9. Recent biophysical data have shown there is a high degree of binding specificity within the extracellular domain between the different family members ^23^. This may explain why we do not see complete loss of all cone bipolars when we delete Pcdh9 as some other family member may act redundantly to prevent cell death. Moreover, recent spatial transcriptomic data from bipolar neurons have revealed subtype identity genes which includes members of the non-clustered Protocadherin family such as Pcdh17 and Pcdh10 are not expressed until later stages of retinal development ^3^. On the other hand, we found Pcdh9 is one of the few non-clustered Protocadherin family members that is expressed at early developmental time points. This may account for why early deletion of full-length of Pcdh9 in Pcdh9 CKO animals result in loss of multiple cone bipolars and not just one particular subtype.

### Mechanisms of cell survival in the developing mouse retina

New emerging data from bipolars and ganglion cells suggest neuronal subtype identity is not determined at birth but arises at later stages of development ^3,39^. This is why it has been difficult to study bipolar subtypes at early developmental time points (P5-13), as most of the antibodies and mouse reporter lines that are bipolar subtype specific are not expressed or do not become subtype-restricted until later stages (>P21). As bipolar subtype identity is not finalized until after synaptogenesis, then there must be some form of plasticity that allows for the appropriate number of bipolars to be specified into each subtype. From our data, we propose bipolar neurons undergo two waves of cell death that ultimately determines the final number of each bipolar subtype. The first wave of bipolar survival relies on full-length Pcdh9 and occurs before synapse formation, whereas the second wave is dependent on homophilic Pcdh9-Pcdh9 interactions of bipolars to their respective synaptic partners. Previous studies in the cancer field have shown Pcdh9 influences cell death by regulating expression of the pro-apoptotic gene, *Bax* ^30^. This is consistent with germline deletion of *Bax^-/-^* that leads to an increase in bipolars of the mouse retina ^14^. Thus, there may be a Pcdh9-mediated mechanism to block the Bax pathway and promote bipolar survival at early stages. However, at later stages Pcdh9 may act in a Bax-independent manner that rely on the extracellular domain forming appropriate homophilic interactions between synaptic partners. This is contrary to prior studies that showed the number of bipolar subtypes are not altered due to changes in photoreceptors or ganglion cells ^14^. However, a caveat of this study was that only OFF cone bipolars (BC2, BC3b, BC4) were analyzed and not ON cone bipolars. Our data revealed ON cone bipolars such as BC7 but not OFF cone bipolars (BC2) were lost due to deletion of Pcdh9 within horizontal cells and amacrine cells, which may account for differences in phenotypes between the two studies. Moreover, OFF cone bipolars and ON cone bipolars form distinct synaptic connections within the outer and inner retina ^1,7^. Dendrites of ON cone bipolars invaginate the cone terminal and form tight associations with the processes of horizontal cells, whereas OFF cone bipolars sit at the base of the cone terminal and do not form invaginating contacts with horizontal cells ^1,7^. This may be the reason why ON cone bipolars are more susceptible to changes in synaptic connectivity with horizontal cells but not OFF cone bipolars.

In conclusion, our work identified a new role for Pcdh9 in bipolar subtype survival during retinal circuit formation. Future studies are needed to determine the downstream mechanisms of how Pcdh9 promotes cell survival and the function of other non-clustered Protocadherins in the developing retina.

### Experimental Procedures

#### Mouse strains

All mouse procedures were approved by the Institutional Animal Care and Use Committee from Baylor College of Medicine and conducted in accordance with NIH guidelines. The floxed allele of Pcdh9 (*Pcdh9^flox/flox^*) and the L385D point mutation (Pcdh9^L385D^) were generated by the Genetically Engineered Rodent Models (GERM) core at Baylor College of Medicine. For the *Pcdh9^flox/flox^* mouse line, LoxP sites were inserted flanking Exon 2 of the *Pcdh9* gene using a CRISPR/Cas9-based knock-in approach as described in ^40^. See **Supplementary Figure 1A**. Briefly, two independent single guide RNAs (sgRNAs) listed in **Table 1** were designed to target the 5’ and 3’ flanking regions of Exon 2. Insertion of the LoxP sites were confirmed by PCR and sanger sequencing as shown in **Supplementary Figure 1B,C** using primers listed in **Table 1**. The L385D point mutation mouse line was generated using a similar CRISPR/Cas9-based knock-in approach ^40^. A sgRNA listed in **Table 1** was used to introduce a single-base pair mutation (ATT>ATC) at 385 position within the extracellular cadherin repeat 4 (EC4). This replaced the original Leucine (L) with Aspartic acid (D) as shown in **Supplementary Figure 2A**. We also introduced an EcoRV restriction enzyme site in frame with the L385D mutation to aid in genotyping. The L385D point mutation was verified by PCR and sanger sequencing (**Supplementary Figure 2B,C**) using primers listed in **Table 1**. In addition, we screened for potential off-targets from the CRISPR editing approach and found no mutations located at these sites (**Table 2**). Both *Pcdh9^flox/flox^* and Pcdh9^L385D^ founders were outcrossed with wild-type C57BL/6J mice for several generations before performing experiments. To conditionally delete Pcdh9 throughout the retina, we used the *Tg(Chx10-EGFP/cre+)* line which has been described previously ^22^. See **Table 3**. To remove Pcdh9 from horizontal cells and amacrine cells, we used the *Tg(Ptf1a-cre)* mouse line described in ^32^ as listed in **Table 3**. Mouse reporter lines such as *Tg(Gus:GFP)* mice were obtained from Jackson laboratories ^41^ and *Tg(Grm6:GFP)* have been previously described in ^2^. See **Table 3**. *Pcdh9^flox/flox^* littermates without cre served as controls for all experiments. Both males and females were used in all experiments.

**Table 1:**
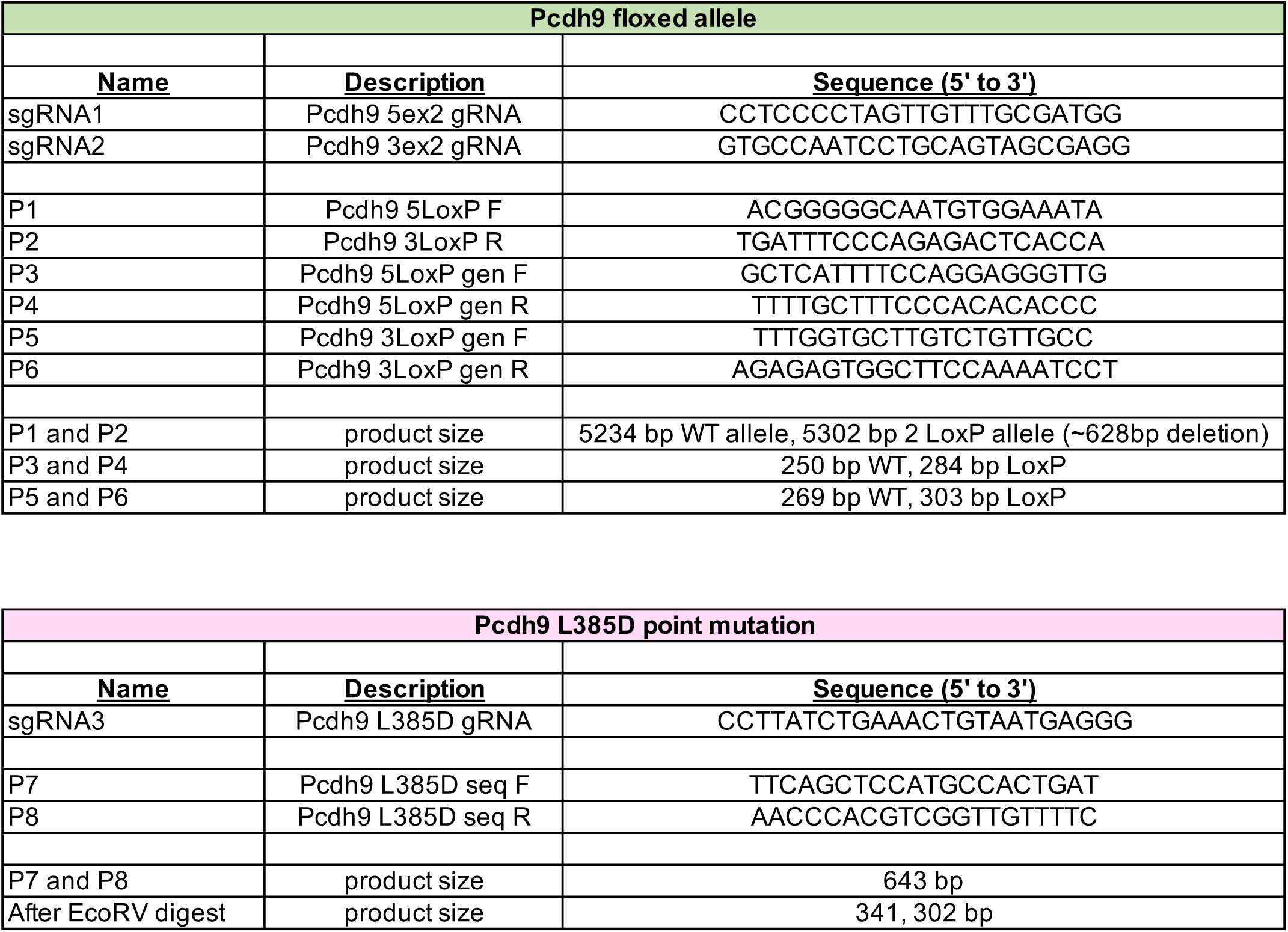
List of primers and single guide RNA (sgRNA) sequences used in this study.

**Table 2:**
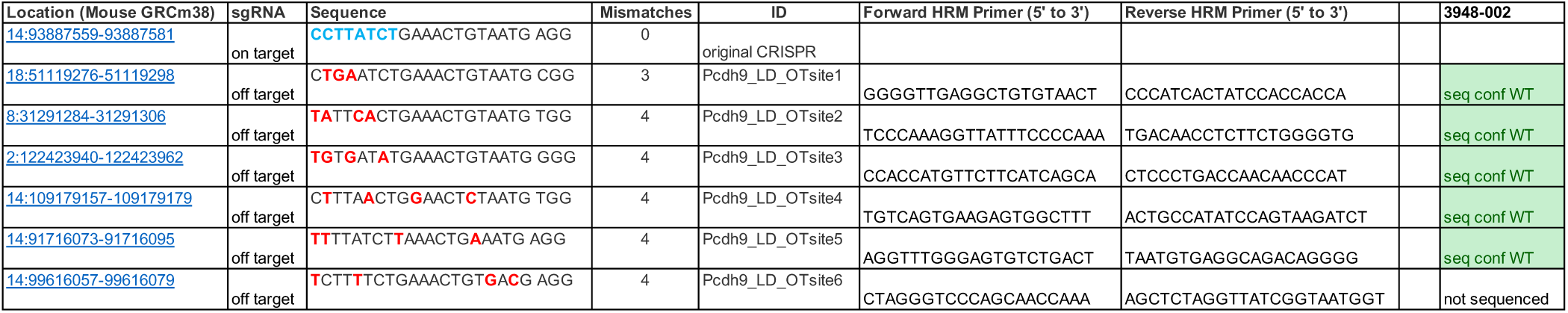
Results from potential off-targets using CRISPR-based approach to generate Pcdh9^L385D^ mice. No mutations were found in the founder line used to propagate the mice used in this study.

**Table 3:**
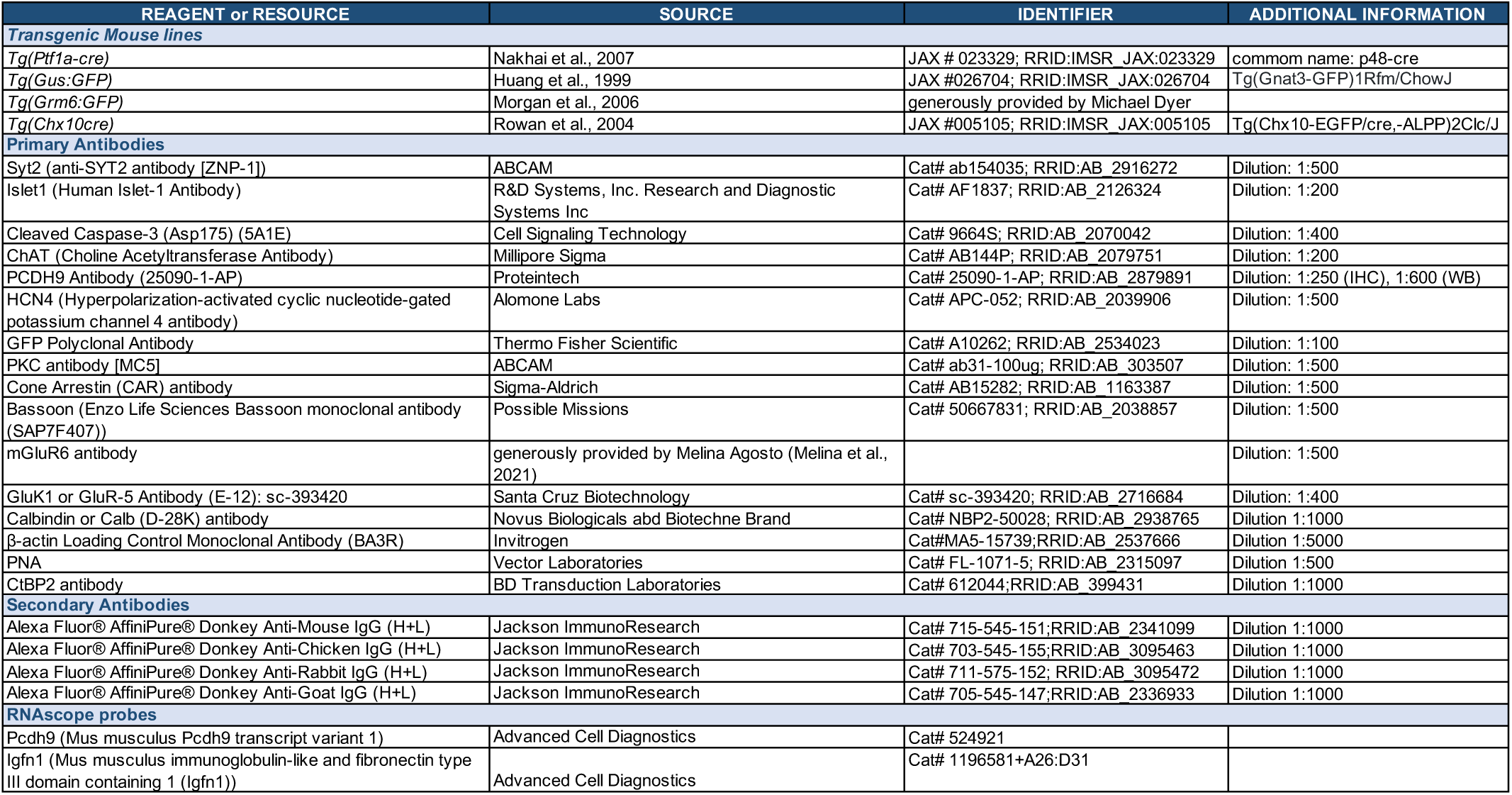
List of reagents used in this study.

#### Immunohistochemistry

Eyes were collected at various developmental time points with P0 designated as the day of birth. Whole eyes were fixed at 60 mins in 4% paraformaldehyde in PBS except for GluK1 staining which were lightly fixed at room temperature for 10 mins. Eye cups were dissected and sectioned at 20µm. Staining was performed as previously described ^29,42^. A list of primary and secondary antibodies used for this study is provided in **Table 3**. Confocal images were acquired on a Zeiss LSM 800 microscope and analyzed using ImageJ version 1.54p and Imaris software version 9.6 (Bitplane, South Windsor, CT, USA).

#### Western blot

Eyes were collected at various developmental time points including P7, P9, P13, and P30. Retinas were dissected in PBS and immediately transferred to ice cold RIPA lysis buffer (Thermo Fisher, Cat#89900) supplemented with 1x protease inhibitor (Roche, Cat#11836170001). Retinal tissue was briefly homogenized by brief sonication on ice. For immunoblotting, 100μg of total protein per sample was loaded on a gel. Proteins were transferred onto a membrane, blocked, and incubated with primary antibodies overnight at 4°C degrees. This was followed by incubation with secondary antibodies for two hours at room temperature. Images were captured using a BioRad gel doc imager.

#### Histological quantification

Data for quantification was collected from 9 different retinal sections from three different animals per group as mentioned in the figure legend. All confocal images were taken from the same central-periphery region of the retina. The *IPLaminator* plugin from ImageJ version 1.54p described in ^24^ was used to measure fluorescence intensity across the IPL with anti-ChAT staining serving as reference for S2 and S4 layers. Quantification of pre- and post-synaptic marker expression was performed using the Spot feature from the Imaris confocal software as previously described in ^29,42^. Puncta with a diameter of 0.6µm was used to detect protein expression of Bassoon. The number of puncta in the OPL per 100μm length was normalized by measuring the total length of the OPL in each individual retinal section. Technical replicates from different retinal sections are shown as small circles and biological replicates from different animals as big circles in all figures. Statistical tests were performed using GraphPad Prism version 9 and details of analysis are provided in the figure legends.

#### RNAscope

Eyes were processed for *in situ* hybridization using RNAscope technology (Advanced Cell Diagnostics). Probe details are provided in **Table 3**. Quantification of *Pcdh9* mRNA and *Igfn1* mRNA puncta was performed using the Imaris confocal software. The number of puncta was normalized to 100μm length of OPL. Details on statistical analysis are provided in the figure legend.

## Acknowledgements

We thank Melanie Samuel, Melinda Agosto, and Michael Dyer for generously providing transgenic mouse lines and antibodies. We are also grateful for Denise Lanza from the GERM core for help with designing the Pcdh9 transgenic mouse lines used in this study. This work was supported by the National Eye Institute (R01EY033037), Research to Prevent Blindness (RPB) Career Development Award to EZS, Center for Alzheimer’s and Neurodegenerative Diseases (CAND) postdoctoral fellowship to MFM, a P30EY002520 and an unrestricted grant from RPB to the Department of Ophthalmology.

## Author Contributions

MFM and EZS designed the experiments. DB, JG, and CCG collected tissue for histological analyses and performed antibody staining. MFM acquired confocal images, performed quantification, and generated figures. MFM and EZS wrote the manuscript.

---

**Supplementary Figure 1:**
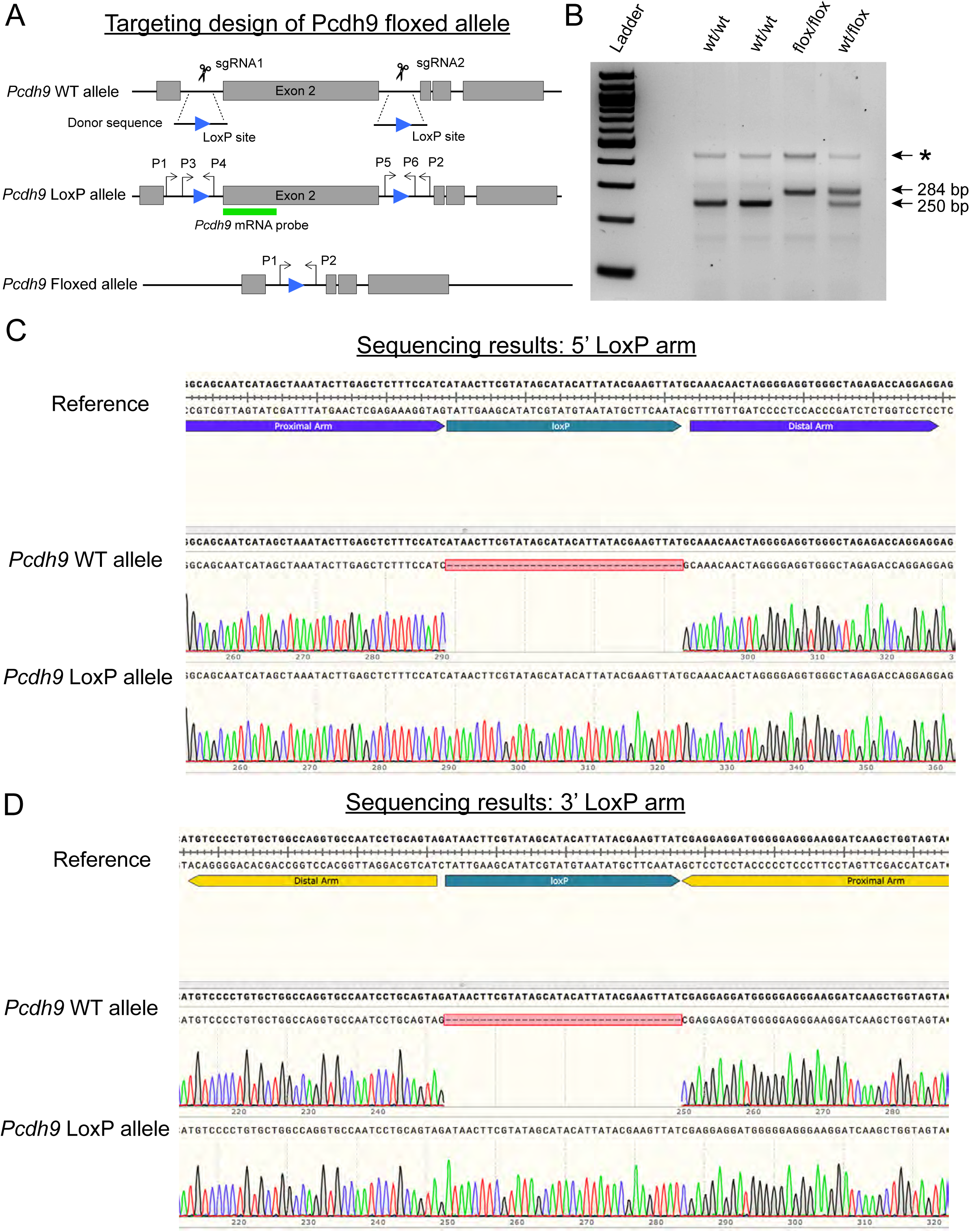
Targeting strategy to generate a floxed allele of Pcdh9 (A-D). Schematic drawing of the targeting strategy using a CRISPR-based approach to insert two LoxP sites flanking Exon 2 of the *Pcdh9* gene in mice (A). Diagnostic gel of the PCR used to detect the 5’ LoxP site using primers P3 and P4 (B). Asterisk denotes a non-specific band that is faintly detected ∼400bp in (B). Sanger sequencing confirms the insertion of the 5’ and 3’ LoxP sites in floxed Pcdh9 mice but not in wild-type Pcdh9 animals (C,D). The reference Pcdh9 sequence contains the LoxP sites and used to confirm insertion of both LoxP sites in floxed Pcdh9 mice in (D).

**Supplementary Figure 2:**
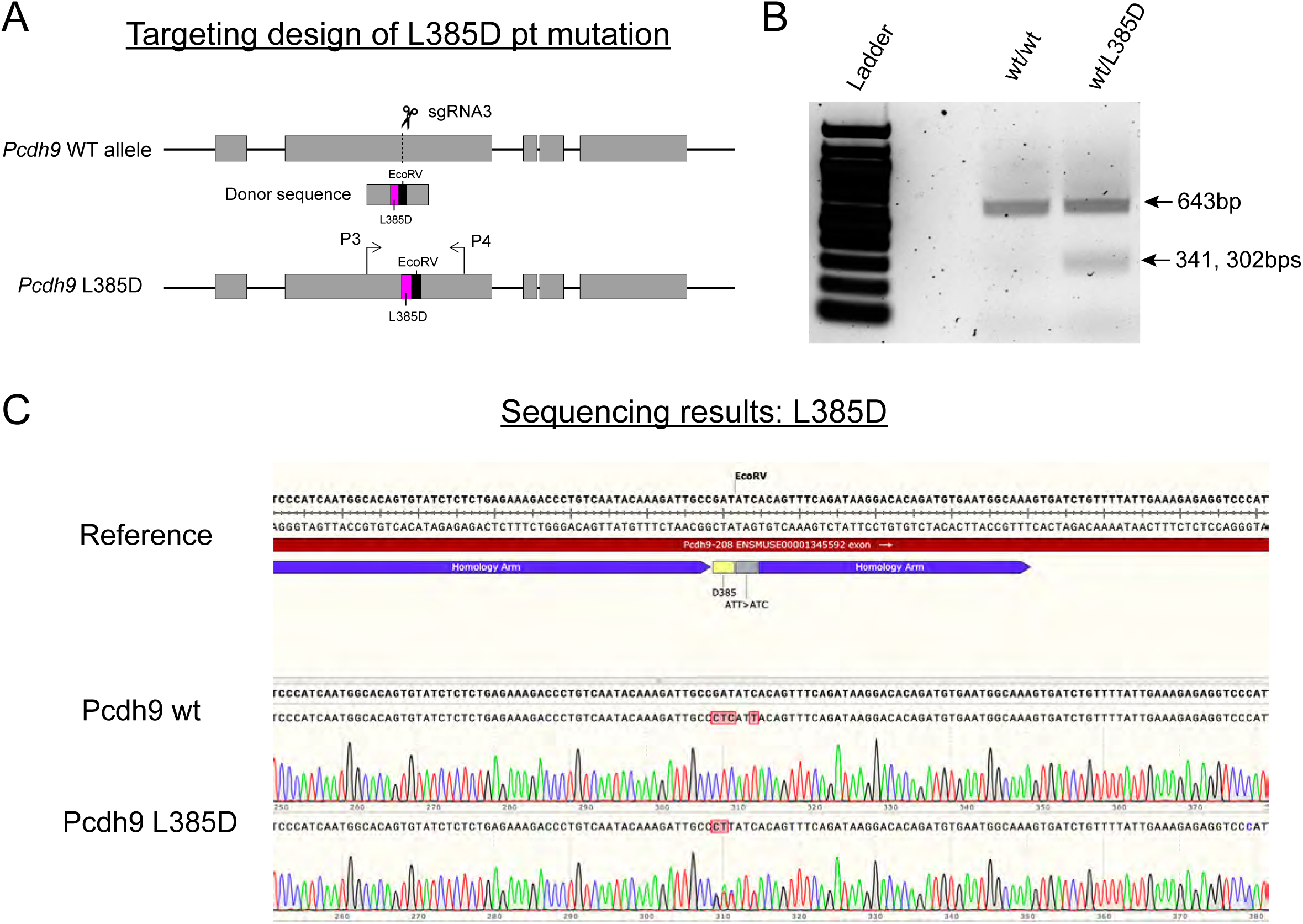
Design strategy to generate L385D point mutation in Pcdh9 (A-C). Schematic drawing of the targeting strategy using a CRISPR-based approach to insert a donor sequence containing the L385D point mutation along with an EcoRV restriction enzyme site (A). Diagnostic gel of the PCR followed by restriction enzyme digest with EcoRV used to detect the presence of the L385D point mutation (B). Wild-type mice show only one undigested band at 643bp, whereas Pcdh9^wt/L385D^ show an additional digested band at 341bp and 302bp (B). Sanger sequencing results confirm the L385D point mutation and the EcoRV site in Pcdh9^L385D^ animals but not in wild-type mice (C).

**Supplementary Figure 3:**
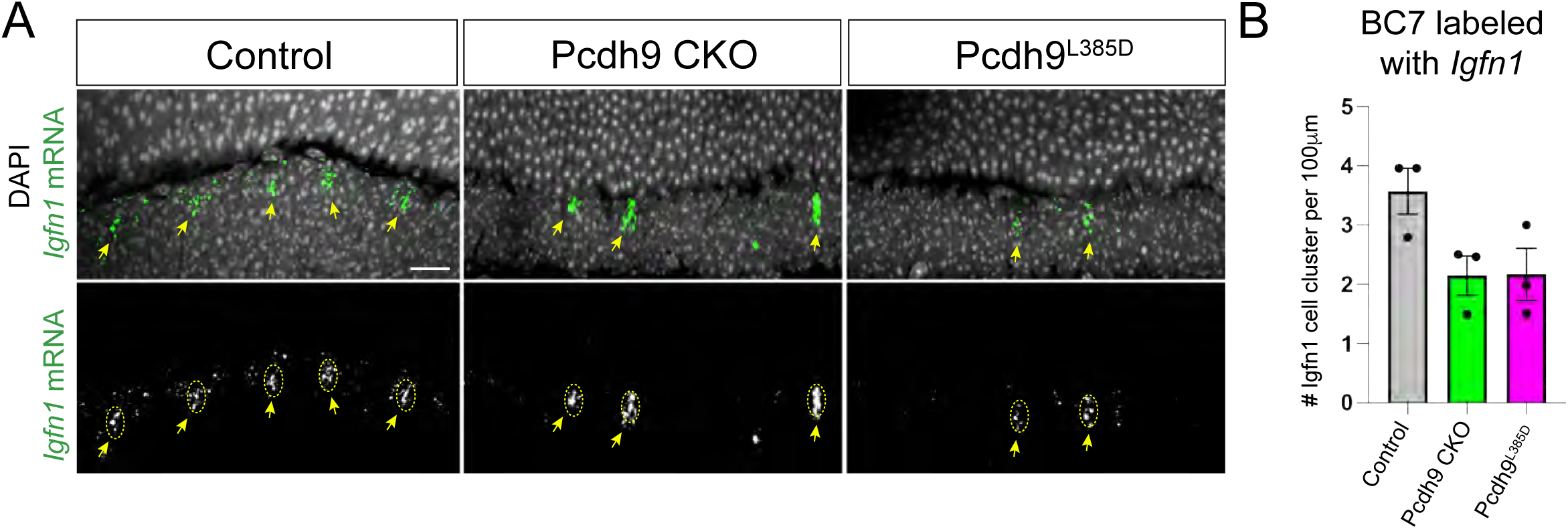
Disruption of Pcdh9 function leads to loss of BC7 neurons (A-B). BC7 neurons were labeled by *in situ* hybridization using the known marker, *Igfn1* (shown in green) in controls, Pcdh9 CKO, and Pcdh9^L385D^ retinas at P30 (A). BC7 neurons are normally evenly spaced across the INL as depicted by yellow dotted circles and arrows as seen in controls in (A). Disruption of Pcdh9 leads to loss of BC7 neurons in Pcdh9 CKO and Pcdh9^L385D^ retinas compared to controls (A). Nuclei were stained with DAPI in (A). Quantification of the total number of *Igfn1*-positive cell clusters (yellow circles) per 100μm OPL length in (A) are shown in (B). Data is represented as mean values ± SEM. Three different retinal sections from at least one animal per group were used for analysis. Scale bar = 25μm.

**Supplementary Figure 4:**
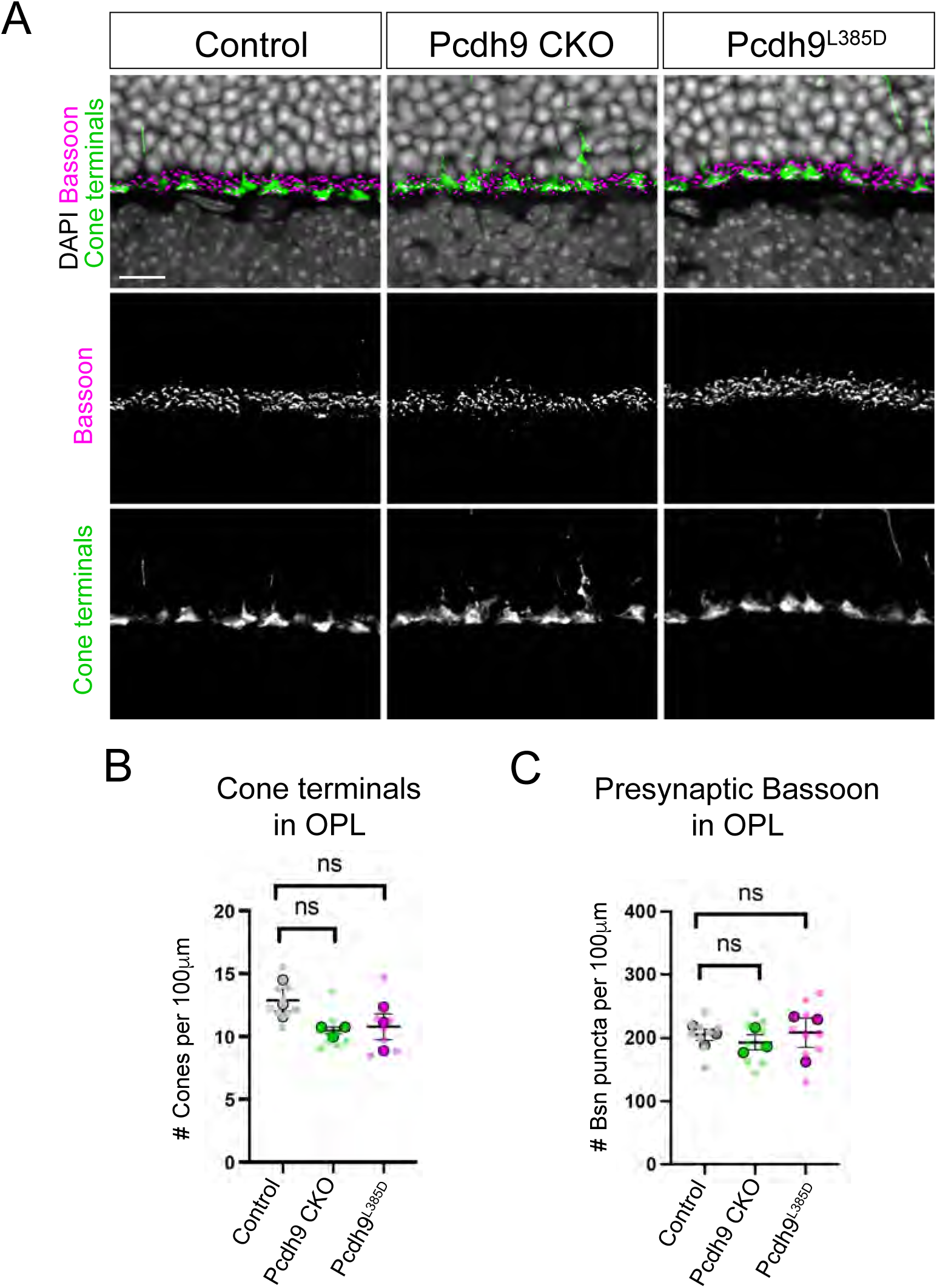
Analysis of cone photoreceptors and pre-synaptic protein expression in Pcdh9 CKO and Pcdh9^L385D^ retinas (A-C). Cone terminals were labeled with anti-CAR (shown in green) in controls, Pcdh9 CKO, and Pcdh9^L385D^ animals at P30 in (A). Retinal sections were also stained for the known pre-synaptic marker, Bassoon (shown in magenta) in (A). The number of cone terminals or pre-synaptic Bassoon protein expression was not statistically different between the three groups (B,C). A total of 9 retinal sections (small circles) from three different animals (big circles) were used for quantification in (B,C). Data is represented as mean values ± SEM and statistical significance determined by an unpaired two-tailed Student’s t test. ns p > 0.05. Scale bar = 25μm.

**Supplementary Figure 5:**
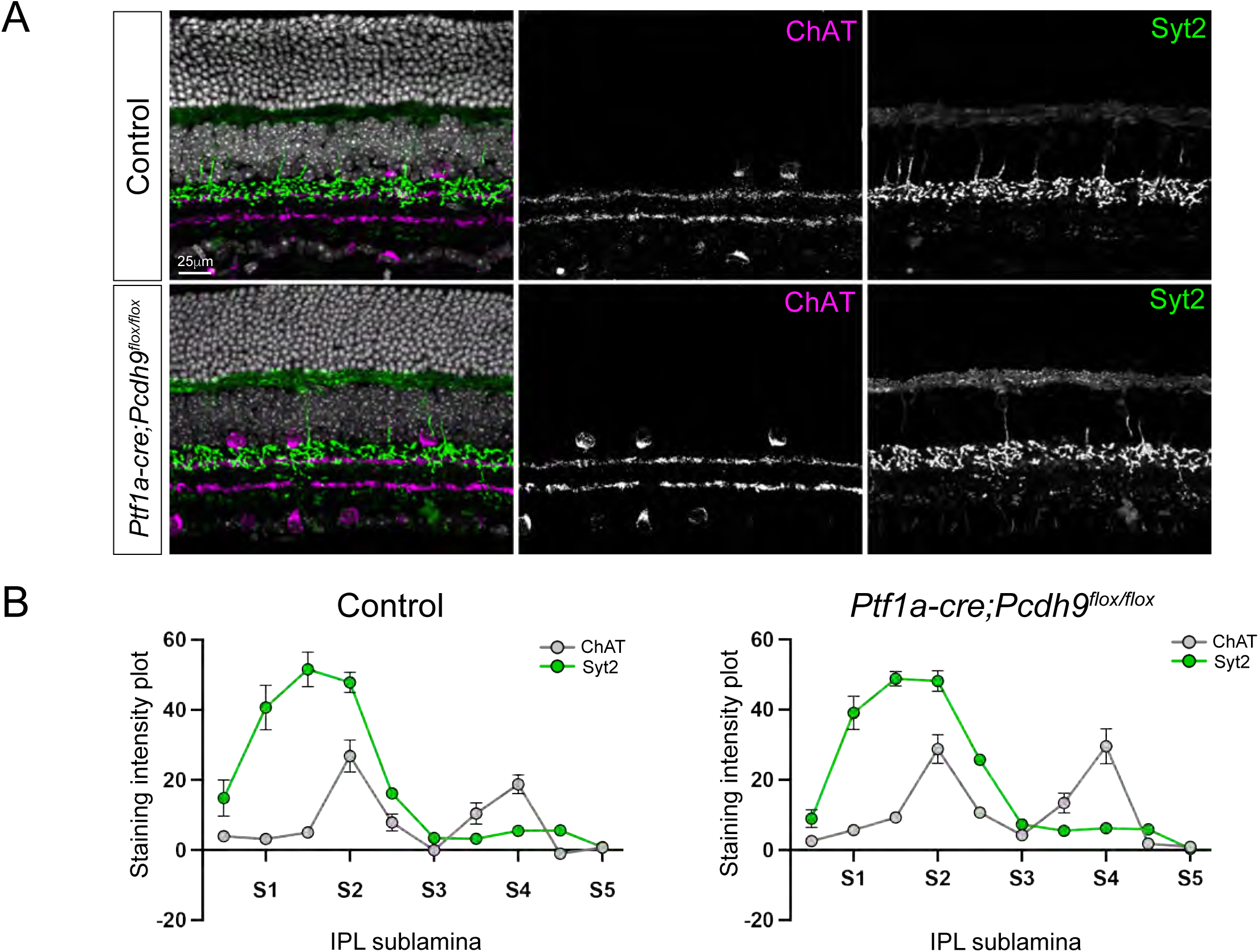
BC2 neurons do not appear affected with deletion of Pcdh9 in horizontal cells and amacrine cells (A-B). BC2 neurons were labeled with anti-Syt2 (green) in controls and *Ptf1a-cre;Pcdh9^flox/flox^* animals (A). Staining with anti-ChAT (magenta) was used as a reference to label S2 and S4 layer of the IPL (A). Nuclei was stained with DAPI in (A). Quantification of the staining intensity across the IPL was performed using the *IPLaminator* tool in controls and *Ptf1a-cre;Pcdh9^flox/flox^* animals as shown in (B). Data are represented as mean values ± SEM. A total of 9 retinal sections from three different animals were used for quantification. Statistical significance was determined by a Holm–Sidak method for multiple comparisons. ns p>0.05. Scale bar on figure.

**Supplementary Figure 6:**
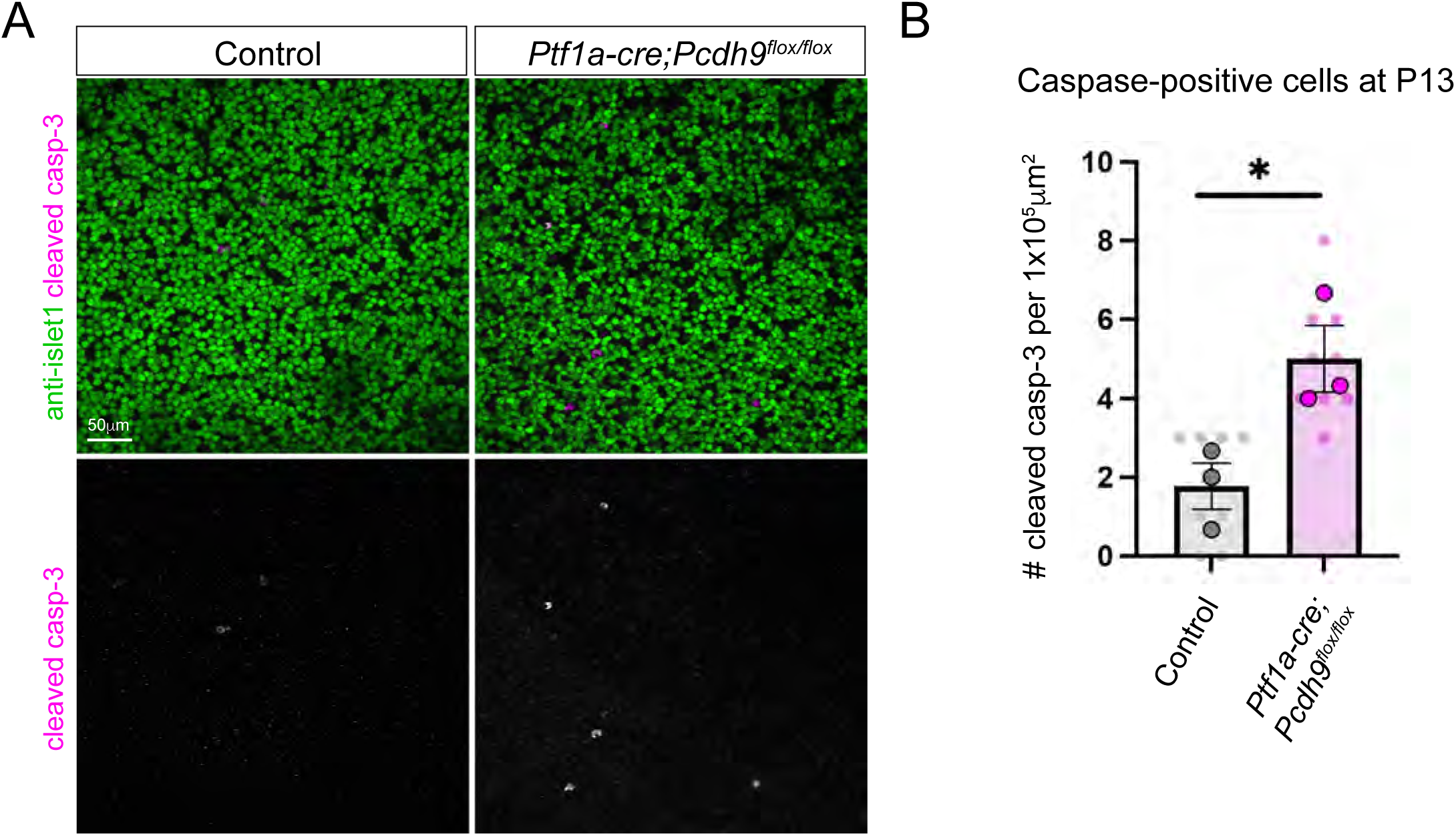
Increased cell death in *Ptf1a-cre;Pcdh9^flox/flox^* compared to controls at P13 (A-B). Dying cells within the bipolar layer were labeled using the antibody against the cleaved form of caspase-3 (cleaved casp-3) shown in magenta and anti-islet1 shown in green (A). Cell counts in controls and *Ptf1a-cre;Pcdh9^flox/flox^* animals are shown in (B). A total of 9 retinal sections (small circles) from three different animals (big circles) were used for quantification. Data are represented as mean values ± SEM with statistical significance determined by an unpaired two-tailed Student’s t test. *p < 0.05. Scale bar = 50μm.

## References

1. Euler, T., Haverkamp, S., Schubert, T., and Baden, T. (2014). Retinal bipolar cells: elementary building blocks of vision. Nat Rev Neurosci 15, 507–519. 10.1038/nrn3783.

2. Morgan, J.L., Dhingra, A., Vardi, N., and Wong, R.O. (2006). Axons and dendrites originate from neuroepithelial-like processes of retinal bipolar cells. Nat Neurosci 9, 85–92. 10.1038/nn1615.

3. West, E.R., Lapan, S.W., Lee, C., Kajderowicz, K.M., Li, X., and Cepko, C.L. (2022). Spatiotemporal patterns of neuronal subtype genesis suggest hierarchical development of retinal diversity. Cell Rep 38, 110191. 10.1016/j.celrep.2021.110191.

4. Sarin, S., Zuniga-Sanchez, E., Kurmangaliyev, Y.Z., Cousins, H., Patel, M., Hernandez, J., Zhang, K.X., Samuel, M.A., Morey, M., Sanes, J.R., and Zipursky, S.L. (2018). Role for Wnt Signaling in Retinal Neuropil Development: Analysis via RNA-Seq and In Vivo Somatic CRISPR Mutagenesis. Neuron 98, 109–126 e108. 10.1016/j.neuron.2018.03.004.

5. Shekhar, K., Lapan, S.W., Whitney, I.E., Tran, N.M., Macosko, E.Z., Kowalczyk, M., Adiconis, X., Levin, J.Z., Nemesh, J., Goldman, M., et al. (2016). Comprehensive Classification of Retinal Bipolar Neurons by Single-Cell Transcriptomics. Cell 166, 1308–1323 e1330. 10.1016/j.cell.2016.07.054.

6. Li, J., Choi, J., Cheng, X., Ma, J., Pema, S., Sanes, J.R., Mardon, G., Frankfort, B.J., Tran, N.M., Li, Y., and Chen, R. (2024). Comprehensive single-cell atlas of the mouse retina. iScience 27, 109916. 10.1016/j.isci.2024.109916.

7. Behrens, C., Schubert, T., Haverkamp, S., Euler, T., and Berens, P. (2016). Connectivity map of bipolar cells and photoreceptors in the mouse retina. Elife 5. 10.7554/eLife.20041.

8. Dunn, F.A., and Wong, R.O. (2012). Diverse strategies engaged in establishing stereotypic wiring patterns among neurons sharing a common input at the visual system’s first synapse. J Neurosci 32, 10306–10317. 10.1523/JNEUROSCI.1581-12.2012.

9. Hellmer, C.B., and Ichinose, T. (2018). Functional and Morphological Analysis of OFF Bipolar Cells. Methods Mol Biol 1753, 217–233. 10.1007/978-1-4939-7720-8_15.

10. Ichinose, T., Fyk-Kolodziej, B., and Cohn, J. (2014). Roles of ON cone bipolar cell subtypes in temporal coding in the mouse retina. J Neurosci 34, 8761–8771. 10.1523/JNEUROSCI.3965-13.2014.

11. Sherry, D.M., Wang, M.M., Bates, J., and Frishman, L.J. (2003). Expression of vesicular glutamate transporter 1 in the mouse retina reveals temporal ordering in development of rod vs. cone and ON vs. OFF circuits. J Comp Neurol 465, 480–498. 10.1002/cne.10838.

12. Anastassov, I.A., Wang, W., and Dunn, F.A. (2019). Synaptogenesis and synaptic protein localization in the postnatal development of rod bipolar cell dendrites in mouse retina. J Comp Neurol 527, 52–66. 10.1002/cne.24251.

13. Young, R.W. (1985). Cell differentiation in the retina of the mouse. Anat Rec 212, 199–205. 10.1002/ar.1092120215.

14. Keeley, P.W., Madsen, N.R., St John, A.J., and Reese, B.E. (2014). Programmed cell death of retinal cone bipolar cells is independent of afferent or target control. Dev Biol 394, 191–196. 10.1016/j.ydbio.2014.08.018.

15. Pequignot, M.O., Provost, A.C., Salle, S., Taupin, P., Sainton, K.M., Marchant, D., Martinou, J.C., Ameisen, J.C., Jais, J.P., and Abitbol, M. (2003). Major role of BAX in apoptosis during retinal development and in establishment of a functional postnatal retina. Dev Dyn 228, 231–238. 10.1002/dvdy.10376.

16. Lee, S.C., Cowgill, E.J., Al-Nabulsi, A., Quinn, E.J., Evans, S.M., and Reese, B.E. (2011). Homotypic regulation of neuronal morphology and connectivity in the mouse retina. J Neurosci 31, 14126–14133. 10.1523/JNEUROSCI.2844-11.2011.

17. West, E.R., and Cepko, C.L. (2022). Development and diversification of bipolar interneurons in the mammalian retina. Dev Biol 481, 30–42. 10.1016/j.ydbio.2021.09.005.

18. Marshall, C.R., Noor, A., Vincent, J.B., Lionel, A.C., Feuk, L., Skaug, J., Shago, M., Moessner, R., Pinto, D., Ren, Y., et al. (2008). Structural variation of chromosomes in autism spectrum disorder. Am J Hum Genet 82, 477–488. 10.1016/j.ajhg.2007.12.009.

19. Bucan, M., Abrahams, B.S., Wang, K., Glessner, J.T., Herman, E.I., Sonnenblick, L.I., Alvarez Retuerto, A.I., Imielinski, M., Hadley, D., Bradfield, J.P., et al. (2009). Genome-wide analyses of exonic copy number variants in a family-based study point to novel autism susceptibility genes. PLoS Genet 5, e1000536. 10.1371/journal.pgen.1000536.

20. Miozzo, F., Murru, L., Maiellano, G., di Iasio, I., Zippo, A.G., Zambrano Avendano, A., Metodieva, V.D., Riccardi, S., D’Aliberti, D., Spinelli, S., et al. (2024). Disruption of the autism-associated Pcdh9 gene leads to transcriptional alterations, synapse overgrowth, and defective network activity in the CA1. J Neurosci 44. 10.1523/JNEUROSCI.0491-24.2024.

21. Brown, S.D., and Moore, M.W. (2012). Towards an encyclopaedia of mammalian gene function: the International Mouse Phenotyping Consortium. Dis Model Mech 5, 289–292. 10.1242/dmm.009878.

22. Rowan, S., and Cepko, C.L. (2004). Genetic analysis of the homeodomain transcription factor Chx10 in the retina using a novel multifunctional BAC transgenic mouse reporter. Dev Biol 271, 388–402. 10.1016/j.ydbio.2004.03.039.

23. Harrison, O.J., Brasch, J., Katsamba, P.S., Ahlsen, G., Noble, A.J., Dan, H., Sampogna, R.V., Potter, C.S., Carragher, B., Honig, B., and Shapiro, L. (2020). Family-wide Structural and Biophysical Analysis of Binding Interactions among Non-clustered delta-Protocadherins. Cell Rep 30, 2655–2671 e2657. 10.1016/j.celrep.2020.02.003.

24. Li, S., Woodfin, M., Long, S.S., and Fuerst, P.G. (2016). IPLaminator: an ImageJ plugin for automated binning and quantification of retinal lamination. BMC Bioinformatics 17, 36. 10.1186/s12859-016-0876-1.

25. Wassle, H., Puller, C., Muller, F., and Haverkamp, S. (2009). Cone contacts, mosaics, and territories of bipolar cells in the mouse retina. J Neurosci 29, 106–117. 10.1523/JNEUROSCI.4442-08.2009.

26. Soto, F., Lin, C.I., Jo, A., Chou, S.Y., Harding, E.G., Ruzycki, P.A., Seabold, G.K., Petralia, R.S., and Kerschensteiner, D. (2025). Molecular mechanism establishing the OFF pathway in vision. Nat Commun 16, 3708. 10.1038/s41467-025-59046-0.

27. Borghuis, B.G., Looger, L.L., Tomita, S., and Demb, J.B. (2014). Kainate receptors mediate signaling in both transient and sustained OFF bipolar cell pathways in mouse retina. J Neurosci 34, 6128–6139. 10.1523/JNEUROSCI.4941-13.2014.

28. Masu, M., Iwakabe, H., Tagawa, Y., Miyoshi, T., Yamashita, M., Fukuda, Y., Sasaki, H., Hiroi, K., Nakamura, Y., Shigemoto, R., and, et al. (1995). Specific deficit of the ON response in visual transmission by targeted disruption of the mGluR6 gene. Cell 80, 757–765. 10.1016/0092-8674(95)90354-2.

29. Pourhoseini, S., Goswami-Sewell, D., and Zuniga-Sanchez, E. (2021). Neurofascin Is a Novel Component of Rod Photoreceptor Synapses in the Outer Retina. Front Neural Circuits 15, 635849. 10.3389/fncir.2021.635849.

30. Wang, C., Tao, B., Li, S., Li, B., Wang, X., Hu, G., Li, W., Yu, Y., Lu, Y., and Liu, J. (2014). Characterizing the role of PCDH9 in the regulation of glioma cell apoptosis and invasion. J Mol Neurosci 52, 250–260. 10.1007/s12031-013-0133-2.

31. Lefebvre, J.L., Kostadinov, D., Chen, W.V., Maniatis, T., and Sanes, J.R. (2012). Protocadherins mediate dendritic self-avoidance in the mammalian nervous system. Nature 488, 517–521. 10.1038/nature11305.

32. Nakhai, H., Sel, S., Favor, J., Mendoza-Torres, L., Paulsen, F., Duncker, G.I., and Schmid, R.M. (2007). Ptf1a is essential for the differentiation of GABAergic and glycinergic amacrine cells and horizontal cells in the mouse retina. Development 134, 1151–1160. 10.1242/dev.02781.

33. Saito, M., Tucker, D.K., Kohlhorst, D., Niessen, C.M., and Kowalczyk, A.P. (2012). Classical and desmosomal cadherins at a glance. J Cell Sci 125, 2547–2552. 10.1242/jcs.066654.

34. Wu, Q., and Maniatis, T. (2000). Large exons encoding multiple ectodomains are a characteristic feature of protocadherin genes. Proc Natl Acad Sci U S A 97, 3124–3129. 10.1073/pnas.97.7.3124.

35. Duan, X., Krishnaswamy, A., De la Huerta, I., and Sanes, J.R. (2014). Type II cadherins guide assembly of a direction-selective retinal circuit. Cell 158, 793–807. 10.1016/j.cell.2014.06.047.

36. Zipursky, S.L., and Sanes, J.R. (2010). Chemoaffinity revisited: dscams, protocadherins, and neural circuit assembly. Cell 143, 343–353. 10.1016/j.cell.2010.10.009.

37. Lefebvre, J.L., Zhang, Y., Meister, M., Wang, X., and Sanes, J.R. (2008). gamma-Protocadherins regulate neuronal survival but are dispensable for circuit formation in retina. Development 135, 4141–4151. 10.1242/dev.027912.

38. Ing-Esteves, S., Kostadinov, D., Marocha, J., Sing, A.D., Joseph, K.S., Laboulaye, M.A., Sanes, J.R., and Lefebvre, J.L. (2018). Combinatorial Effects of Alpha- and Gamma-Protocadherins on Neuronal Survival and Dendritic Self-Avoidance. J Neurosci 38, 2713–2729. 10.1523/JNEUROSCI.3035-17.2018.

39. Shekhar, K., Whitney, I.E., Butrus, S., Peng, Y.R., and Sanes, J.R. (2022). Diversification of multipotential postmitotic mouse retinal ganglion cell precursors into discrete types. Elife 11. 10.7554/eLife.73809.

40. Lanza, D.G., Gaspero, A., Lorenzo, I., Liao, L., Zheng, P., Wang, Y., Deng, Y., Cheng, C., Zhang, C., Seavitt, J.R., et al. (2018). Comparative analysis of single-stranded DNA donors to generate conditional null mouse alleles. BMC Biol 16, 69. 10.1186/s12915-018-0529-0.

41. Huang, L., Shanker, Y.G., Dubauskaite, J., Zheng, J.Z., Yan, W., Rosenzweig, S., Spielman, A.I., Max, M., and Margolskee, R.F. (1999). Ggamma13 colocalizes with gustducin in taste receptor cells and mediates IP3 responses to bitter denatonium. Nat Neurosci 2, 1055–1062. 10.1038/15981.

42. Goswami-Sewell, D., Bagnetto, C., Gomez, C.C., Anderson, J.T., Maheshwari, A., and Zuniga-Sanchez, E. (2023). betaII-Spectrin Is Required for Synaptic Positioning during Retinal Development. J Neurosci 43, 5277–5289. 10.1523/JNEUROSCI.0063-23.2023.

